# Fibrinogen anchors for micropatterning of active proteins and subcellular receptor relocalisation

**DOI:** 10.1101/2020.09.04.256875

**Authors:** Joseph L. Watson, Samya Aich, Benjamí Oller Salvia, Andrew A. Drabek, Stephen C. Blacklow, Jason Chin, Emmanuel Derivery

## Abstract

Protein micropatterning allows proteins to be precisely deposited onto a substrate of choice, and is now routinely used in cell biology and in vitro reconstitution. However, a drawback of current technology is that micropatterning efficiency can be variable between proteins, and that proteins may lose activity on the micropatterns. Here, we describe a general method to enable micropatterning of virtually any protein at high specificity and homogeneity while maintaining its activity. Our method is based on an anchor that micropatterns well, Fibrinogen, which we functionalized to bind to common purification tags. This enhances micropatterning on various substrates, facilitates multiplexed micropatterning, and dramatically improves the on-pattern activity of fragile proteins like molecular motors. Furthermore, it enhances the micropatterning of hard to micropattern cells. Last, this method enables subcellular micropatterning, whereby complex micropatterns simultaneously control cell shape and the distribution of transmembrane receptors within that cell. Altogether, these results open new avenues for cell biology.

## Introduction

Micropatterning, also known as molecular printing, is the process by which molecules are precisely deposited onto a substrate of choice with micrometer resolution. Micropatterning is now routinely used in all area of biomedical research, for example in DNA microarrays, which have been around for decades and rely on surfaces being printed one spot at a time^1^. Decades of photolithography technology led to the advent of parallelized techniques for the affordable and straightforward manufacturing of protein micropatterns on a variety of substrates^2,3^. This led to breakthroughs in biology, allowing researchers, for example, to print cell adhesion proteins to constrain cell shape and thus reveal the physical basis of cell polarity^4^ and spindle orientation during mitosis in cultured cells^5^. Concomitantly, the ability to print purified proteins onto glass brought better control in *in vitro* reconstitution studies, which strengthened our understanding of the dynamics of cytoskeleton contractility^6^, as well as the physiology of cytoskeletal polymers^7,8^. Finally, it was recently demonstrated that electron microscopy grids could be micropatterned with cell adhesion molecules^9,10^. The ability, therefore, to constrain the position and shape of cells on grids is poised to solve the decade-old problem that cells adhere much more efficiently to electron-impermeable grid bars than electron-permeable carbon mesh.

Subtractive patterning by photolithography offers an efficient and convenient method to generate protein micropatterns on any substrate. In this technique, the substrate is first uniformly coated with an antifouling agent such as PEG^3^ that blocks non-specific protein binding. This homogenous coat is then precisely etched by photoscission using a deep UV light source and a quartz photomask^11^ to allow specific adsorption of the protein of interest. While broadly adopted, this method has the inconvenient feature that each time one wants to make a novel pattern, a new mask has to be manufactured, adding cost and time to the scientific process. As noted early on with photobleaching-based techniques^12,13^, using a microscope instead of a mask to project the illumination pattern has the major advantage that spatial light modulators, such as DMDs (Digital Micromirror Devices) can be used to shape the illumination pattern at will. In addition, using a microscope *de facto* enables multiplexed patterning of multiple proteins^13^, because the micropatterned protein can be imaged and the subsequent illumination pattern precisely aligned with it. Multi-protein patterning is much more tedious to achieve with masks or protein stamps due to difficulties in alignment^14^. Importantly, while deep UV is not compatible with glass-based microscope optics, Fink and colleages^15^ found that micropatterning of PEG-coated coverslips could be achieved with regular UV light with a benzophenone photosensitizer. Later elegant work^16^ by Strale and colleagues integrated all these developments (microscope/DMD/photosensitizer/UV) into a pioneering technique they termed LIMAP (light-induced molecular adsorption), which they demonstrated allows convenient multiprotein patterning in a commercial microscope.

While micropatterning offers tremendous possibilities for biomedical research, a potential limitation of the technology is the highly variable patterning efficiency amongst different proteins. In particular, patterning selectivity (how much protein adsorbs to the patterned compared to the non-patterned region) and patterning homogeneity (how the adsorption density varies within the pattern) vary greatly between proteins (see also Fig. 1 and S4 of this paper). For instance, if a given cell type requires a key extracellular protein to adhere to the substrate, but that protein does not micropattern efficiently, this cell type currently cannot be used in micropatterning assays. This variability between proteins is even more problematic when micropatterning multiple proteins, because it implies that the different proteins will not necessarily be patterned at a consistent density. For complex micropatterning experiments, such as testing the effects of opposing gradients of signalling molecules, variable and inhomogeneous adsorption efficiencies is thus a huge challenge. Furthermore, while LIMAP-mediated multiplexed micropatterning offers unprecedented avenues for biology, it may bring additional problems. In particular, because of the sequential nature of the patterning process, nonspecific binding between successive patterns is possible, where the second protein binds to the first pattern if it has not been completely quenched (a phenomenon we refer to as cross-adsorption in this paper). In addition, the buffers and Reactive Oxygen Species (ROS) needed for LIMAP patterning may not be optimal for maintaining the activity of proteins, so while a protein might pattern well, it might have lost its activity by the time the user starts the experiment. Another general source of activity loss when micropatterning is that the direct adsorption of a protein onto the surface may induce unfolding. All these limitations may either discourage researchers from doing such complex experiments, or force them to perform long and painstaking optimizations of their patterning process.

**Figure 1.**
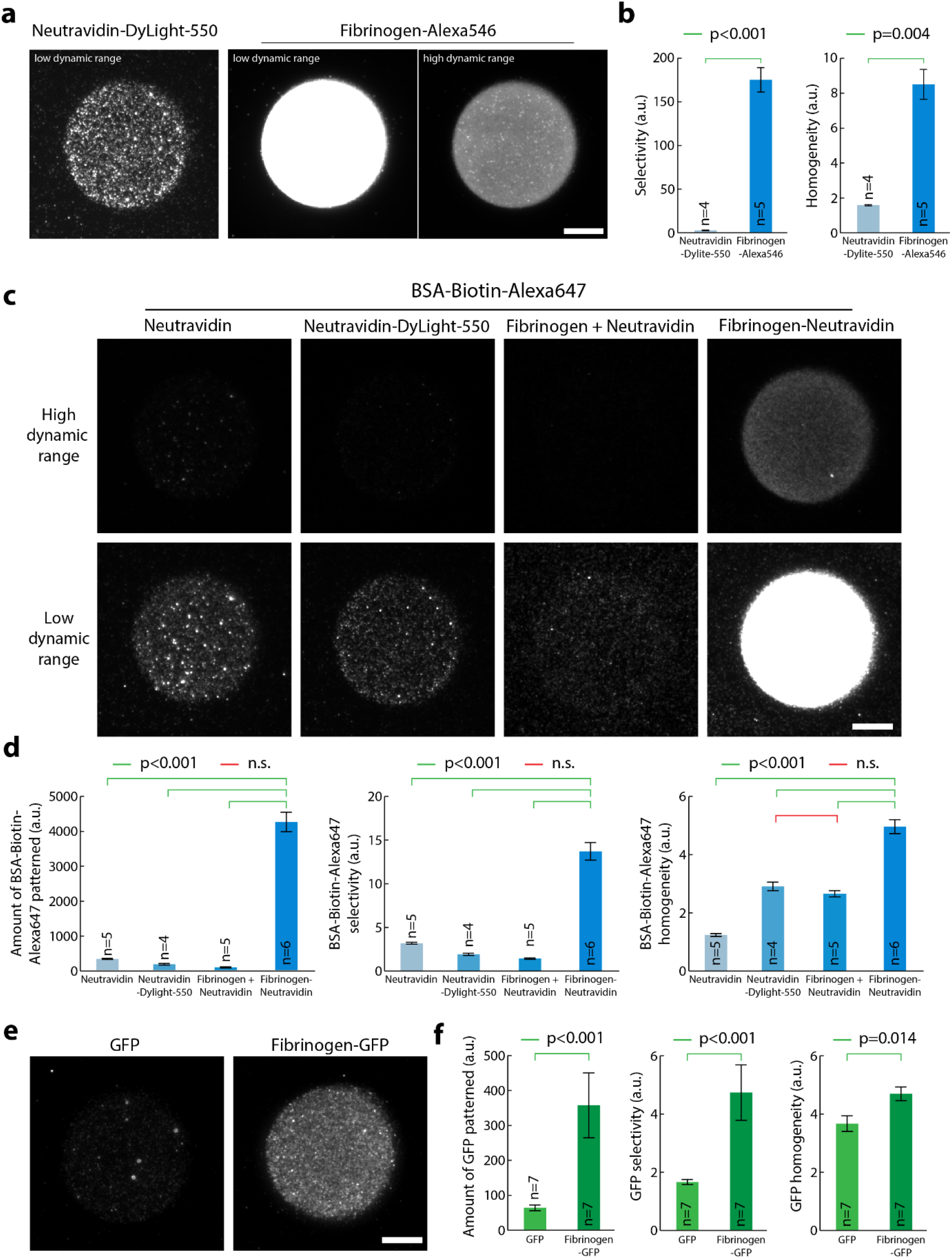
Fibrinogen anchors improves selectivity and homogeneity of micropatterns on PLL-PEG surfaces. (a) Fibrinogen-Alexa546 (50 μg/ml) and NeutrAvidin-Dylight-550 (50 μg/ml) were micropatterned on PLL-PEG-coated glass using LIMAP with identical UV exposure and pattern. After washing, red fluorescence of the patterns was imaged by TIRFM using identical settings (left and middle panels). Alternatively, a five-fold lower exposure was used (right panel). (b) Quantification of the effects seen in (a) Mean ± SEM of the selectivity and homogeneity (see methods). Statistics were performed using a Mann-Whitney Rank Sum Test, n: number of patterns measured. Fibrinogen quantitatively patterns better than NeutrAvidin. (c) NeutrAvidin, NeutrAvidin-DyLight-550, NeutrAvidin mixed with Fibrinogen or NeutrAvidin fused to Fibrinogen (Fibrinogen-NeutrAvidin) were micropatterned on PLL-PEG-coated glass using LIMAP with identical UV exposure, pattern shape and protein concentration (50 μg/ml). After pattern quenching and washing, BSA-Biotin-Alexa647 (5 μg/ml) was added for 5 min, then the sample was washed and BSA-Biotin-Alexa647 fluorescence was imaged by TIRFM. Two different dynamic ranges for visualisation were used for each sample (top versus bottom line) so that each lane could be represented with the same dynamic range. (d) Quantification of the amount of protein bound to the pattern (left panel), pattern selectivity (middle panel) and homogeneity (right panel) in the sample presented in (c). Statistics were performed using an ANOVA1 test followed by a Tukey post-hoc test after log10 transformation of the data (p<0.001). Fibrinogen-NeutrAvidin enhances significantly the selectivity and the homogeneity of BSA-Biotin-Alexa647 patterns. (e) GFP and Fibrinogen GFP were micropatterned on PLL-PEG-coated glass using LIMAP with identical UV exposure, pattern shape and protein concentration (50 μg/ml). After washing, GFP fluorescence was imaged by TIRFM. (f) Quantification of the amount of protein selectively bound to the pattern (left panel, as well as pattern selectivity (middle panel) and homogeneity (right panel) in the sample presented in (e). Statistics were performed using a Student’s t-test (after log10 transformation of the data for left and middle panels). The selectivity and homogeneity of Fibrinogen GFP patterns is significantly better than that of GFP alone. Statistics were performed using a Mann-Whitney Rank Sum Test, n: number of patterns measured. Scale bars: 10 μm.

Here, we describe a general method to enable micropatterning of virtually any protein at a high specificity and homogeneity. Our technology relies on a protein anchor that patterns well, Fibrinogen, which is functionalized to recognize tags commonly used in protein purification, or that can be added to already-purified proteins. We show that this can quantitatively improve patterning of hard-to-micropattern proteins on various substrates. In particular, we demonstrate robust patterning of Concanavalin A (ConA), which thereby enhances the micropatterning of hard-to-micropattern cells, such as *Drosophila* cells. Furthermore, we show that our technology provides an advantage for LIMAP-mediated multiprotein patterning as by design, not only does it allow for all proteins to be micropatterned with similar homogeneity and low non-specific binding, but it also shields proteins of interest from any harm induced by the micropatterning process because they are bound to the micropattern after said process. We also demonstrate that our technology allows the control of the subcellular localization of membrane receptors by simultaneously micropatterning cell adhesion proteins and receptor ligands at high density. Altogether, we hope that our contribution will facilitate micropatterning experiments and open new avenues for biological research.

## Results

We started by comparing the micropatterning efficiency of two proteins broadly used for micropatterning, NeutrAvidin^7,8,17^ and Fibrinogen^11,18,19^. Except for figure 4, all micropatterns in this study were obtained following the LIMAP protocol^16^ using a UV-projector in a fluorescence microscope and the photosensitizer 4-benzoylbenzyl)trimethylammonium bromide (BBTB), see Fig. S1 and methods. To quantitatively compare patterns throughout this paper, we evaluated both pattern selectivity (that is, how much protein adsorbs to the patterned compared to the non-patterned region) and pattern homogeneity (that is, how the adsorption density varies within the pattern) [see also methods]. As seen in Fig.1 a-b both the specificity and the homogeneity of Fibrinogen micropatterns are quantitatively higher than that of NeutrAvidin micropatterns. As micropatterning is an adsorption process, its efficiency is expected to vary according to the buffer composition according to changes in protein folding and surface charges. Accordingly, we found that Fibrinogen patterning is quantitatively improved in low-salt carbonate buffer compared to PBS buffer (Fig. S2), consistent with the known propensity of Fibrinogen to precipitate in the presence of salts. For this reason, all patterns in this paper were generated in carbonate buffer (see methods).

We thought to exploit the inherent high micropatterning efficiency of Fibrinogen to improve the micropatterning of hard-to-micropattern proteins, by conjugating them together. Fibrinogen would thus act as the “anchoring” moiety, ensuring that both micropatterning selectivity and homogeneity are high and reproducible, while the protein of interest would provide the “specificity” part, bringing the function required for the experiment. This would also potentially improve accessibility, and thereby activity, of the micropatterned protein, as not all of it would not be facing the glass (which may occur with direct patterning due to potential preferred orientation of protein deposition). Constraining protein orientation would also mitigate potential surface-induced protein unfolding. Fibrinogen is an ideal anchoring moiety for this as it is commercially available in gram-scales, is easy to work with, and on the contrary to other commonly available proteins like BSA, precipitates natively with very low amounts of Ammonium Sulfate, a property we could exploit to facilitate the separation of functionalized products (see methods). We thus established robust protocols for the biochemical functionalization of Fibrinogen, allowing it to bind to virtually any protein of interest, as well as to commonly used purification tags.

The conjugation method relied on the modification of Fibrinogen exposed amines with a heterobifunctional crosslinker that enabled conjugation with the target protein in a 2- or 3-step procedure (see Fig. S3 and methods). In particular, we derived Fibrinogen-GFP, Fibrinogen-ConA (a lectin that binds to insect cells^20^), Fibrinogen-NeutrAvidin (Fig. S3a) to bind to biotinylated targets, Fibrinogen-GBP to bind to GFP-tagged proteins (Fig. S3b GBP stands for GFP-binding Peptide, a nanobody against GFP^21^) and Fibrinogen-biotin (Fig. S3c) to bind to streptavidin/NeutrAvidin fusions, as well as to biotinylated targets using a NeutrAvidin sandwich (Fibrinogen-biotin/NeutrAvidin/biotinylated-protein of interest). Importantly, since Fibrinogen contains numerous exposed amines, it can be functionalized with multiple molecules. For instance, biotin and ATTO490LS can be combined (Fibrinogen-Biotin-ATTO490LS) to micropattern biotinylated target while at the same time facilitating pattern alignment by fluorescence microscopy (see below). Note that these conjugation protocols are general and apply to virtually any protein of interest. Furthermore, these optimized protocols are straightforward, use commercially available chemicals and do not require any specialized biochemistry equipment such as an FPLC.

As a proof of concept, we compared the micropatterning efficiency of a biotinylated target, BSA-Biotin-Alexa647 through either NeutrAvidin or Fibrinogen-NeutrAvidin patterns. This is a more relevant situation than just comparing the patterning efficiency of Fibrinogen-NeutrAvidin versus NeutrAvidin, as what matters for experiments is the density of micropatterned biotin-binding sites, which is not necessarily the same as the density of micropatterned NeutrAvidin. As it is always the same fluorescent protein being micropatterned, in addition to determining the pattern selectivity and homogeneity, we also estimated the amount of protein being specifically bound to the pattern from the fluorescence intensity. As seen in Fig. 1c-d, Fibrinogen-NeutrAvidin provides a four-fold improvement of the selectivity and homogeneity of micropatterned biotinylated targets, as well as the 13-fold improvement of amount of protein being specifically patterned. This was true for all Fibrinogens, see for instance Fibrinogen-GFP offering higher selectivity, homogeneity and amount of patterned protein than GFP alone (Fig. 1e, see also f for quantification). Importantly, while all experiments described until now were performed on PLL-PEG-coated glass, the Fibrinogen toolkit described here also improves micropatterning on other commonly used substrates, such as PEG-silane^7,8,17^ (Fig. S4).

We then thought to apply our technology to multiprotein-patterning. A key advantage of using the LIMAP protocol within a microscope^16^ is that it enables straightforward multiprotein patterning as the user can both image one micropattern and, in the same instrument, readily align a second pattern with respect to this pattern (*ad infinitum*). As mentioned above, this technology has three potential issues: i) lack of homogeneity between patterns due to the high variability of patterning between proteins; ii) cross-adsorption between successive protein applications (aka successive patterns) because of incomplete quenching and iii) potential activity loss due to inappropriate buffers and/or ROS generation. An additional issue is that while being able to image a micropattern in the microscope to align the next one is a key advantage, this requires the first micropatterned protein to be fluorescent. However, fluorescent derivatives of proteins are not necessarily as active as their unlabelled counterparts: for instance, we found that fluorescent Neutravidin (NeutrAvidin-Dylight-550), while micropatterning fine on Fibrinogen-biotin, display a more than 6-fold reduction in its ability to bind to biotinylated targets compared to unlabelled NeutrAvidin (Fig. S4). Doping the unlabelled protein to micropattern with some other fluorescent protein as a tracer is not a solution either, as, as seen in Fig. 1, micropatterning efficiencies widely differ between proteins, so the fluorescent tracer will not necessarily be a reliable proxy for the unlabelled protein to be micropatterned.

The Fibrinogen toolkit described here could address all these limitations at once. Indeed, first, it would ensure that patterning homogeneity is similar for all patterned proteins, as it is always the same protein anchor that is patterned (Fibrinogen). Second, Fibrinogen anchors would simplify quenching steps after the micropatterning process, as again it is always the same anchor that will have to be quenched. Third, Fibrinogen anchors functionalized against common purification tags would enable one to first micropattern all the anchors, then add the proteins of interest in their buffer of choice as the last step of the sample preparation just before the actual experiment starts, therefore ensuring that the activity of the proteins of interest is conserved (Fig. 2a). Last, each non-fluorescent Fibrinogen anchor can be doped with trace amounts of fluorescent Fibrinogen for pattern alignment purposes without risking these trace amounts to affect the patterning efficiency of the Fibrinogen anchor. This last issue can also be alleviated by fluorescently labelling Fibrinogen anchors (see Fig. 3, 5, 6).

**Figure 2.**
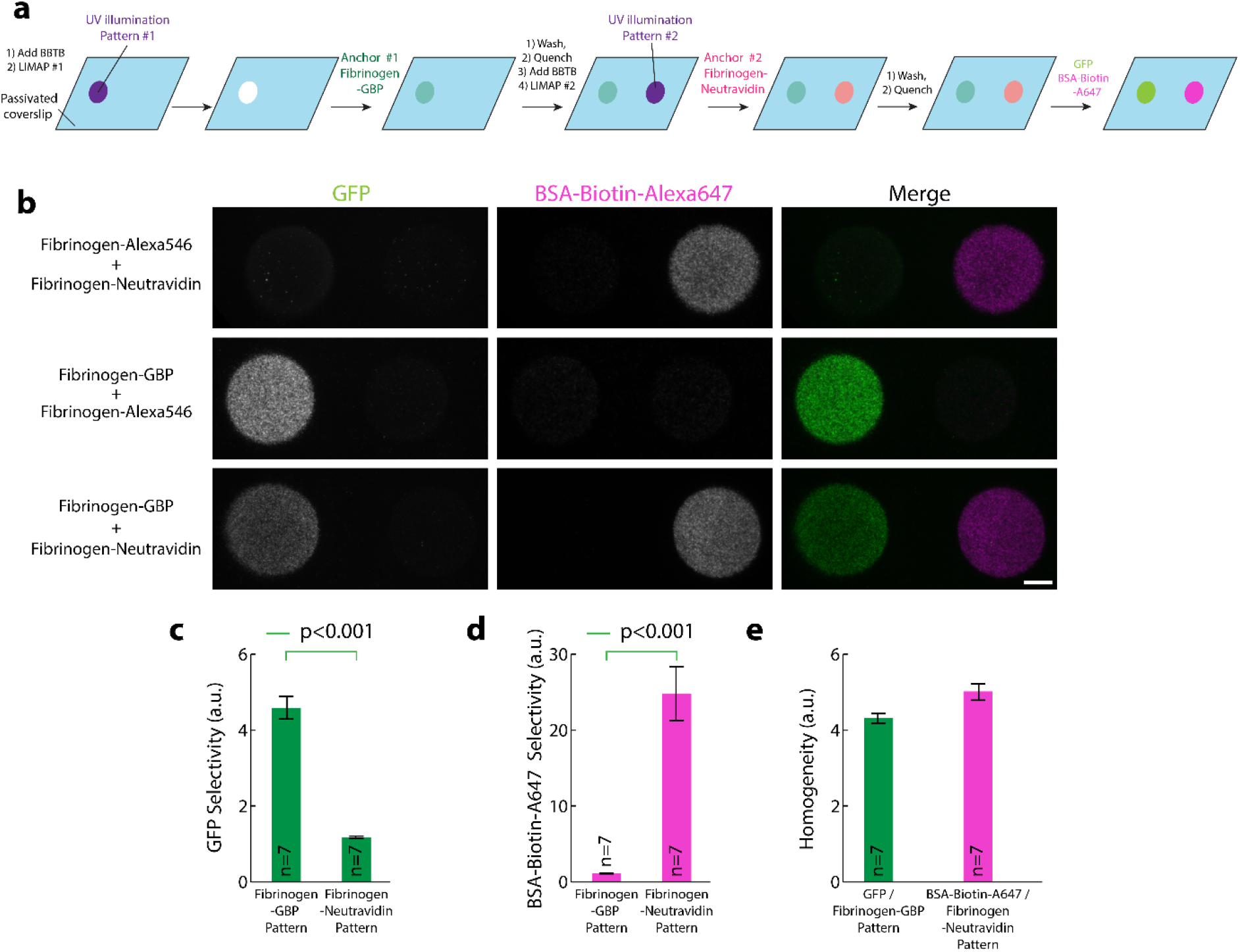
Fibrinogen anchors facilitate multiplexed micropatterning of proteins. (a) Scheme illustrating the different steps for multiplexed micropatterning using Fibrinogen anchors. Note that the two proteins to be micropatterned (GFP and BSA-Biotin-Alexa647) are added together after the patterning process. This implies that their activity can be maintained because they can be kept in their optimal buffer and are not exposed to UV-induced ROS, nor the BBTB. (b) Bottom line: multiplexed patterning of GFP and BSA-Biotin-Alexa647 (5 μg/ml) with Fibrinogen-GBP and Fibrinogen-NeutrAvidin anchors (100 μg/ml) using the scheme depicted in (a) onto PLL-PEG-coated glass. Top and middle lines: controls of bottom panel by exchanging Fibrinogen-GBP by Fibrinogen-Alexa546 (top line) or by exchanging Fibrinogen-NeutrAvidin by Fibrinogen-Alexa546 (middle line) followed by co-injection of GFP and BSA-Biotin-Alexa647. Note that there is high specificity of the protein of interests for their respective anchor and minimum overlap between the two proteins. This implies that the successive patterns of Fibrinogen anchors have been efficiently quenched (otherwise the proteins of interest could bind to the wrong pattern). (c) Quantification of the increased specificity of GFP for the Fibrinogen-GBP anchor compared to the Fibrinogen-NeutrAvidin anchor. (d) Quantification of the increased specificity of BSA-Biotin-Alexa647 for the Fibrinogen-NeutrAvidin anchor compared to the Fibrinogen-GBP anchor. (e) Similarity of the patterning homogeneity of GFP onto Fibrinogen-GBP patterns and BSA-Biotin-Alexa647 on Fibrinogen-NeutrAvidin patterns. Statistics in (c) and (d) were performed using a Mann-Whitney Rank Sum Test. n: number of patterns measured. Scale bar: 10 μm.

As a proof of concept, we patterned two proteins, GFP and Biotin-BSA-Alexa647 using Fibrinogen-GBP and Fibrinogen-NeutrAvidin anchors, respectively (Fig. 2a). As seen in Fig. 2b, Fibrinogen anchors enable reliable micropatterning of these two proteins, with quantitatively minimal crosstalk between the two channels (see Fig. 2c-d for quantitative comparison of patterning selectivity of the two proteins for the two anchors). In addition, pattern homogeneity was similar for both proteins (Fig. 2e), suggesting that both proteins were patterned at a consistent density. We found that multiplexed patterning was extremely reproducible with this protocol, and found that the major source of variability between samples rather came from occasional drying of the coverslip during the patterning process (Fig. S5).

As mentioned above, it is not enough that a protein micropatterns well, its activity must also be maintained on the pattern. Loss of activity during patterning can have multiple origins: the patterning process itself can be detrimental to proteins due to ROS generation and buffer composition when doing LIMAP, but also, more generally, since patterning is an adsorption-based process, proteins will pattern with their “stickier” side facing the glass, which could be the face harbouring the active site, therefore affecting activity or inducing unfolding. Fibrinogen anchors alleviate the buffer/ROS issue, but they potentially could also improve orientation and conformational freedom as the protein of interest is not itself patterned but rather attached to the pattern via a flexible linker.

We thought to test this hypothesis using a commonly-used assay relying on adsorbed proteins, namely the Microtubule (MT) gliding assay^22^ (Fig. 3). In this assay, kinesin molecular motors are adsorbed onto glass, and motors with their motor domain facing away from the glass can then move MTs around. While a biotinylated fragment comprising the motor domain and the dimerising coiled-coil of *Drosophila* kinesin 1^23^ (noted Kin1-biotin) is efficiently micropatterned onto PLL-PEG substrates (Fig. 3a-d), its activity is dramatically reduced when doing so: microtubules move slowly (Fig. 3c, j for quantification, and see also Movie S1) and frequently pause during their motion (Fig. 3c, k for quantification). Conversely, micropatterning the same Kin1-biotin through a “sandwich” on top of Fibrinogen-Biotin-ATTO490LS and NeutrAvidin (Fig. 3e-h), restored expected gliding speeds (Fig. 3g, j for quantification) with very rare pauses (Fig. 3g, k for quantification, and see also Movie S1). Importantly, in all these experiments, we only detected motion in the patterned regions (Fig. 3i), suggesting that MTs bound to the surface outside the pattern were not bound by kinesin but rather by non-specific interactions with the surface. This demonstrates that Fibrinogen anchors are a reliable way to ensure high activity of micropatterned proteins, most likely by ensuring a high density of correctly-orientated proteins, and by mitigating surface-induced and ROS-induced denaturation of the micropatterned proteins of interest.

We then decided to apply our Fibrinogen technology to facilitate applications that would otherwise be hard with existing micropatterning techniques. Insect cells like *Drosophila* S2 cells require lectins such as Concanavalin A (ConA) to adhere on glass^20^. However, ConA does not micropattern well in our hands (Fig. 4a, compare “pattern” channel with Fibrinogen for the small patterns on the right-hand side of each image), thereby limiting the possibility of adhering these cells to small micropatterns, and therefore to micropattern single S2 cells. Note that this is mainly a problem for small micropatterns, as cells manage to adhere partially to large micropatterns (Fig. 4a, left-hand side of all images). We thus derived Fibrinogen-ConA (Fig. S2a), which dramatically improves micropatterning efficiency of S2 cells, even onto small, single cell, micropatterns (Fig. 4a, compare “merge” channel between ConA and Fibrinogen-ConA, see also Fig. 4b-c for quantification of the cell density and specificity on the small micropatterns). The improvement in adhesion was even clearer with medium-sized micropatterns, to which several cells adhered per pattern (Fig. S6).

**Figure 3.**
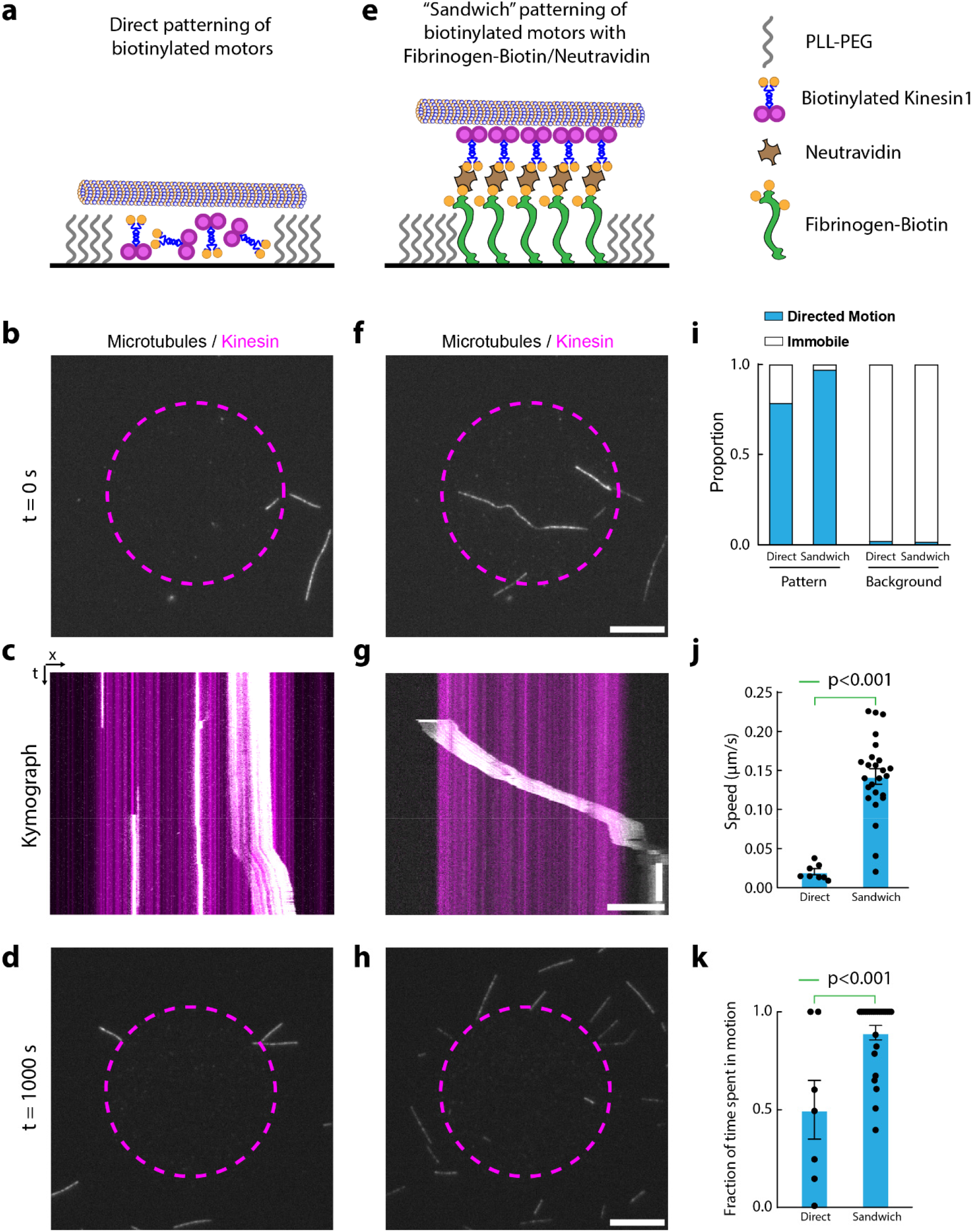
Fibrinogen anchors facilitate micropatterning of active motors. Biotinylated-Kinesin1 motors (Kin1-Biotin) were micropatterned on PLL-PEG-coated glass using LIMAP either directly (a-d) or indirectly through a Fibrinogen-Biotin-ATTO490LS/NeutrAvidin sandwich (e-h), see methods. After washing and quenching, GMPCPP-stabilized fluorescent microtubules were added in the presence of ATP and their motion observed by TIRFM. Dashed purple lines delineate the kinesin pattern as imaged either through post-labelling of Kin1-Biotin by streptavidin-Alexa647 for direct patterning (a-d) or by Fibrinogen-Biotin-ATTO490LS fluorescence for the indirect labelling (e-h). (i) Quantification of the proportion of motile MT in all conditions reveals that MTs outside the pattern are immobile, in contrast to MTs landing inside the pattern (n = 14/34/47/112 respectively). (j-k) Quantification of the speed (j) and fraction of time spent moving processively (k) reveals that MTs move faster and with less pauses on Fibrinogen-biotin-mediated Kin1-Biotin micropatterns (directly patterned n = 7, sandwich n = 25). This suggest that indirect micropatterning of Kin1-biotin through Fibrinogen-biotin ensures high activity of the motor on the pattern. Statistics in (j) and (k) were performed using unpaired t tests. n: number of microtubules. Scale bars: 10 μm (b, d, f, h) and 10 μm/ 1 min (c, g).

**Figure 4.**
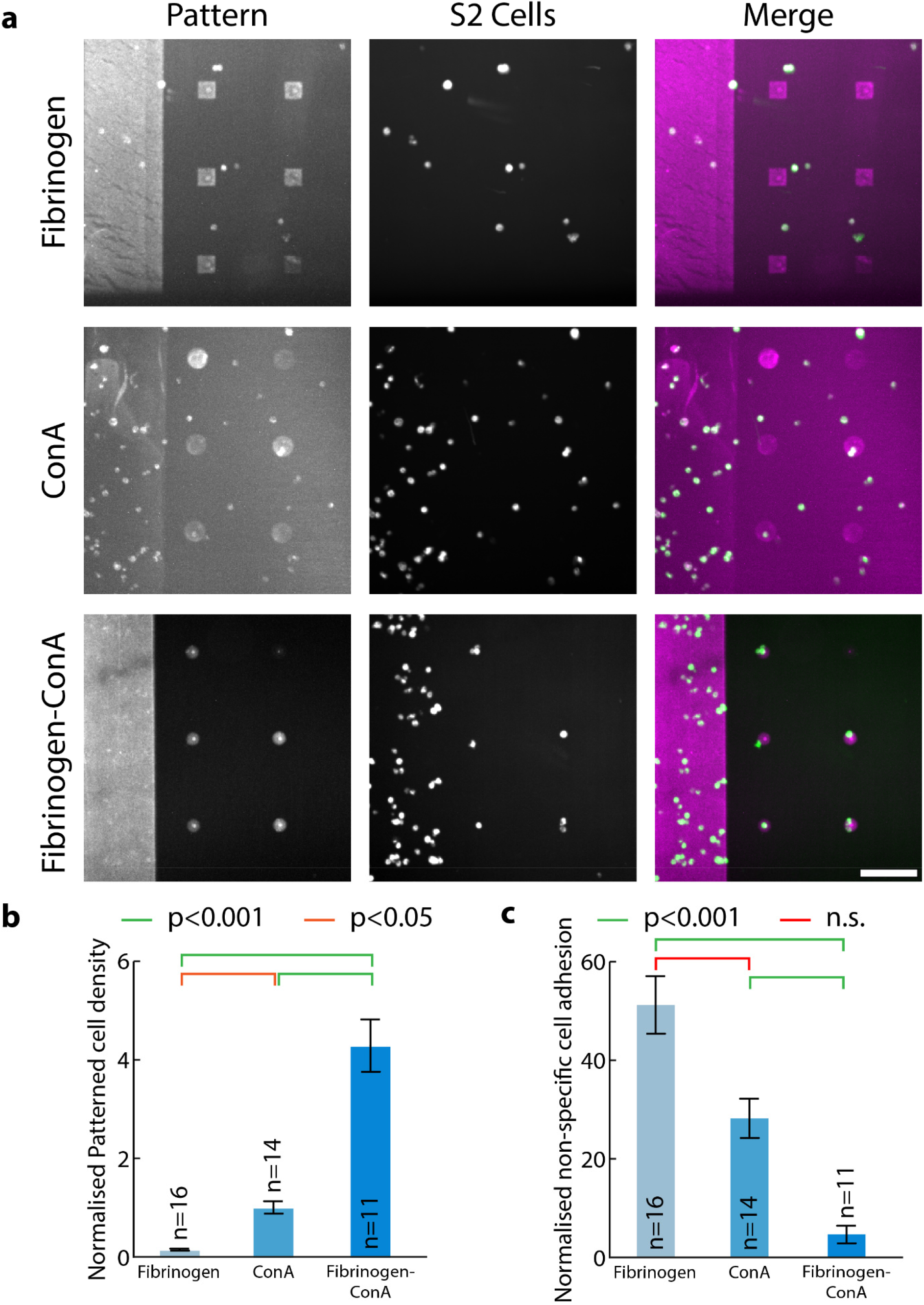
Fibrinogen anchors enahnces the micropatterning of hard-to-pattern cells. (a) Fibrinogen (doped with 10% Fibrinogen-Alexa546), ConA (doped with 10% Rhodamine ConA) or Fibrinogen-ConA (doped with 10% Fibrinogen-Alexa546) were micropatterned at 50 μg/ml onto PLL-PEG-coated glass using deep-UV and a chromium mask. Coverslips were washed, and S2 cells were added for 1h, before addition of SiR tubulin to label cells, for 30 minutes. After washing, cells and micropatterns were imaged by spinning disc confocal microscopy. While S2 cells do manage to adhere on larger ConA patterns, micropatterning efficiency of ConA is lower than that of Fibrinogen (compare left panels), and therefore Fibrinogen-ConA enables micropatterning of S2 cells onto small, single cell, patterns. (b-c) Quantification of the effects seen in A (Fibrinogen n = 16, ConA n = 14, Fibrinogen-ConA n = 11). Normalized patterned cell density (b) is significantly higher on small Fibrinogen-ConA patterns compared to ConA or Fibrinogen patterns, while the normalized non-specific adhesion (c) is higher for Fibrinogen and ConA and Fibrinogen-ConA (see methods for details). Statistics in (b) were performed using an ANOVA1 test followed by a Tukey post-hoc test (p<0.001) while statistics in (c) were performed using a Kruskal-Wallis test. n: number of fields of view analysed. Scale bar: 100 μm.

We then thought to combine all the advantages of our technology to open a new avenue in micropatterning by achieving *subcellular* micropatterning, whereby the position of proteins within cells can be imprinted from the outside via a micropattern. To achieve this, we made dual micropatterns with one pattern to anchor the cell, and a second pattern to relocalise a transmembrane receptor via an active ligand presented in the right orientation (Fig. 5a, 6a, 6d). Achieving this quantitatively obviously relies on the ability to achieve multiplexed micropatterning of proteins while maintaining their activity and proper orientation. As a proof of concept, we used NIH/3T3 cells stably expressing a model receptor composed of a transmembrane segment fused to an extracellular GBP and an intracellular mScarlet (GBP-TM-mScarlet), and dual-micropatterns composed of a Fibrinogen/Fibronectin anchoring pattern, and a Fibrinogen-Biotin-ATTO490LS/Streptavidin-GFP-GFP relocalising pattern. Streptavidin-GFP-GFP refers to a fusion between streptavidin and two copies of GFP. Importantly, this setup achieved specific and quantitative relocalisation of the GBP-TM-mScarlet construct onto the GFP micropattern (Fig. 5b, c). Protein relocalisation happened in a matter of minutes when cells entered in contact with the GFP-positive part of the micropattern during spreading, and was then stable over time (Fig. 5d,e and Movie S2). Importantly, this result showcases the advantages of the high micropatterning densities and modularity that are achievable with our Fibrinogen toolbox, as micropatterns offering lower GFP densities, such as direct patterning of GFP or Fibrinogen-GFP with a low degree of labelling, did not induce noticeable GBP-TM-mScarlet relocalisation (Fig. S7a-b).

**Figure 5.**
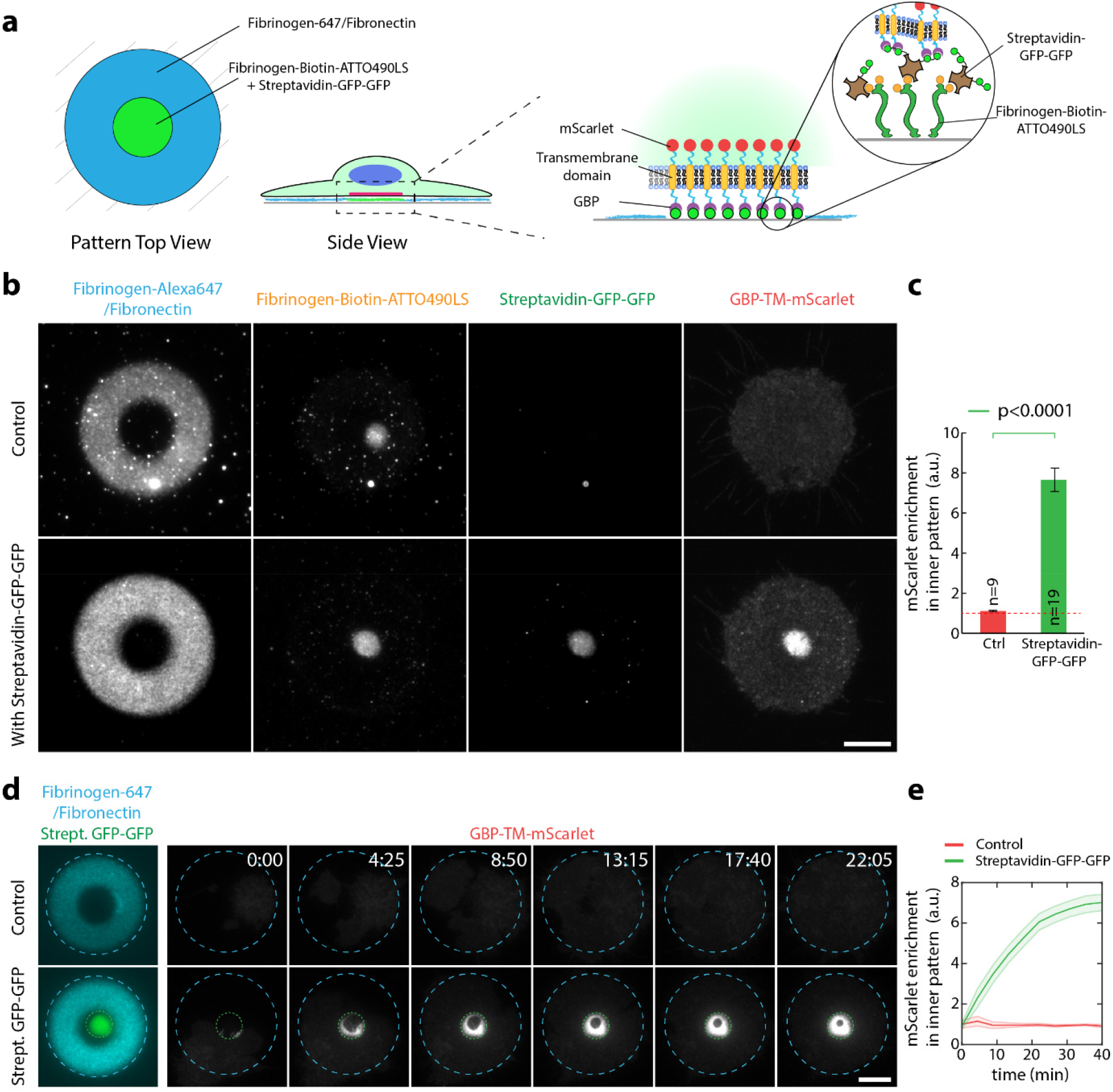
Fibrinogen anchors allow the subcellular micropatterning of a synthetic receptor fused to a cortical protein of interest. (a) Experimental scheme: NIH/3T3 cells stably and constitutively expressing GBP-TM-mScarlet were allowed to spread on dual patterns of Fibronectin/Fibrinogen-Alexa647 and Fibrinogen-Biotin-ATTO490LS/streptavidin-GFP-GFP, and were then imaged live by TIRF microscopy. (b) Efficient relocalisation of the GBP-TM-mScarlet construct onto an area defined by the extracellular pattern in live cells. (c) Quantification of the effects seen in (b). (d) Cells as in (b) were imaged by TIRF microscopy during spreading to evaluate the kinetics of GBP-TM-mScarlet recruitment onto the GFP micropattern. Fibrinogen and GFP micropatterns are figured in blue and green dashed lines, respectively. (e) quantification of the effects seen in (d). Statistics were performed using a Student’s t-test. n: number of cells analysed. Scale bar: 10 μm.

**Figure 6.**
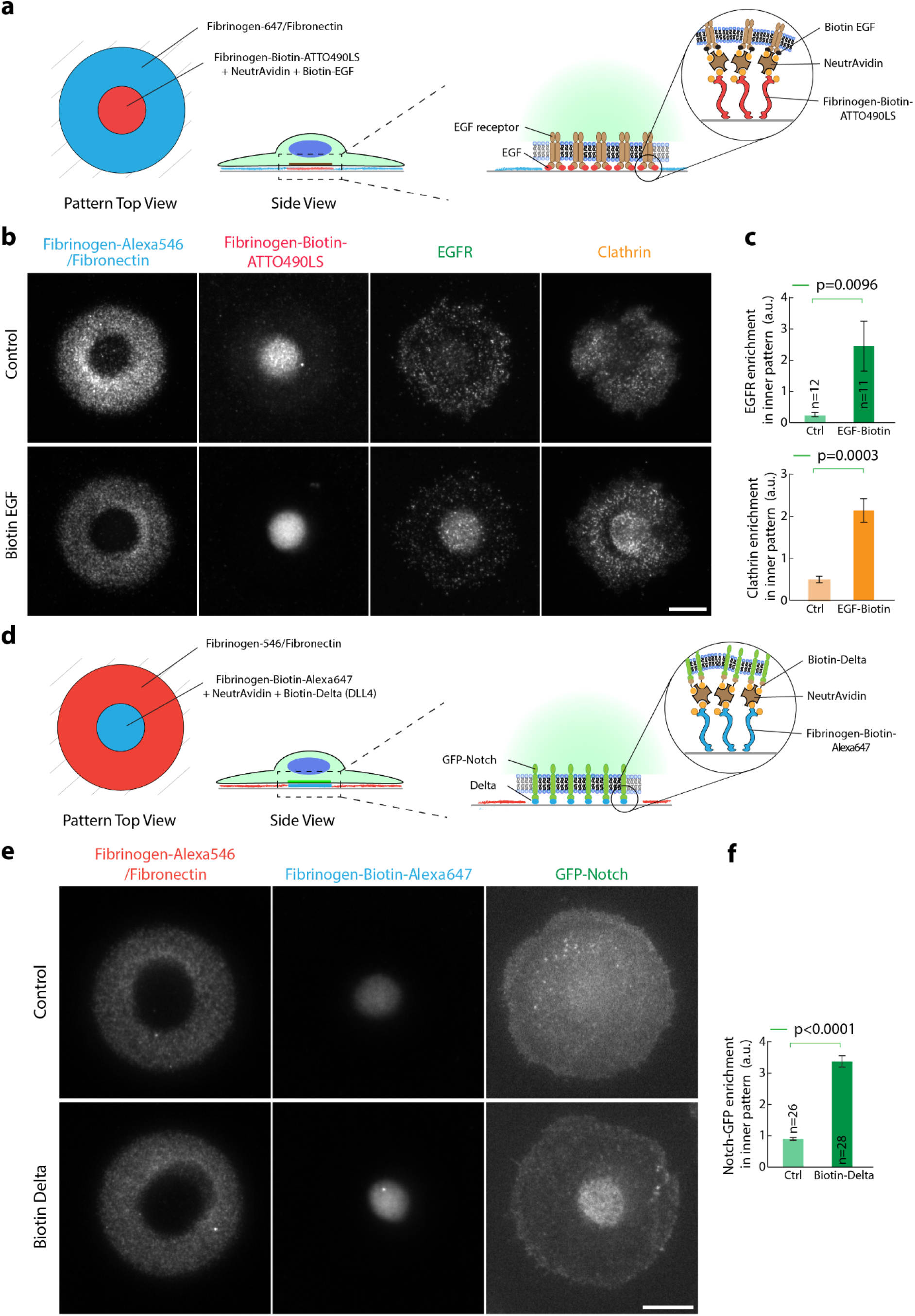
Fibrinogen anchors allow the subcellular micropatterning of endogenous receptors. (a) Experimental scheme: Serum-starved HeLa cells were allowed to spread on dual patterns of Fibronectin/Fibrinogen-Alexa546 and Fibrinogen-Biotin-ATTO490LS/NeutrAvidin/biotin-EGF, then fixed and processed for immunofluorescence against EGFR and Clathrin. (b) Efficient relocalisation of the EGFR onto the active EGF micropattern. Note that Clathrin is also recruited onto the pattern. (c) Quantification of the effects seen in (b). (d) Experimental scheme: U2OS cells stably expressing GFP-Notch1 were allowed to spread on dual patterns of Fibronectin/Fibrinogen-Alexa647 and Fibrinogen-Biotin-ATTO490LS/NeutrAvidin/biotin-DLL4, and imaged live 20 minutes later by TIRF microscopy. (e) Efficient relocalisation of GFP-Notch1 onto the active DLL4 micropattern. (f) Quantification of the effects seen in (e). Statistics were performed using a Student’s t-test. n: number of cells analysed. Scale bar: 10 μm.

We then generalized this concept to endogenous receptor/ligand pairs to establish controlled signalling platforms at the surface of cells. First, a pattern of a soluble ligand, namely the epidermal growth factor (EGF), proved efficient at relocalising its cognate receptor (EGFR), when anchored via a Fibrinogen-biotin/NeutrAvidin sandwich (Fig. 6a, b, see also c for quantification). This was not observed without the addition of biotinylated-EGF (Fig. 6b,c).Importantly, the poor direct micropatterning of fluorescent EGF (Fig. S7c) made this dual-patterning assay impossible without Fibrinogen anchoring. Conversely, the modular nature of the fibrinogen-anchoring technology means that very high densities of active EGF could be achieved using a neutravidin/Fibrinogen-biotin amplification strategy, akin to those used to increase GBP-TM-mScarlet relocalisation (Fig 5a,b). Interestingly, with EGFR relocalisation, we found that Clathrin was also relocalised to the EGF/EGFR pattern (Fig. 6b, see also c for quantification), suggesting that the EGF micropattern traps EGFR in an intermediate along the endocytic pathway, as expected if the pattern activates the physiological signalling cascade downstream of EGFR activation but prevents EGFR endocytosis because of the strong link to the pattern. Similarly, a pattern of a ligand normally exposed at the surface of cells, Delta, proved efficient at relocalising its cognate receptor, GFP-Notch, in live cells (Fig. 6d-f). As before with the synthetic GFP/GBP interaction, relocalisation of GFP-Notch by micropatterned Delta occurred in the time scale of minutes as cells spread onto the pattern, and remained stable over time (Fig. S8 and movie S3).

## Discussion

The technology developed in this paper allows the micropatterning of virtually any protein of interest with high selectivity and homogeneity. We envision that this will open new avenues for cell biology applications, for instance to micropattern cell types that could not be micropatterned before due to the lack of patterning efficiency of their extracellular matrix of choice. In particular, the enhanced micropatterning of *Drosophila* cells which we achieve in this work (Fig. 4) opens a new avenue for cell biology as it will allow us, through the isolation of primary *Drosophila* cells, to combine quantitative geometrically-defined cell culture with the genetic tractability of *Drosophila*. Importantly, while we focused here on UV-micropatterning in a microscope^16^, our technology is general and applies to any light-induced micropatterning technique, such as Deep UV-Quartz-mask based techniques for instance (See Fig. 4)^11^. The ability to functionalize Fibrinogen with nanobodies could also be a benefit for the clustering of receptors against which antibodies have been developed due to the elegant work by Pleiner and colleagues who recently released an extensive toolkit of nanobodies targeting nearly all commonly used primary antibodies^24^. Indeed, these nanobodies are compatible with their rational cysteine-based nanobody engineering protocol^25^, which we rely on for Fibrinogen functionalization with GBP. Importantly, while we focused here on a few commonly used protein tags, there is no reason why other binders/ligands could not be fused to Fibrinogen, such as BG (to bind to SNAP-tagged proteins), Strep-Tactin^26^ (to bind to Strep-Tag-tagged proteins), amylose (to bind to Maltose Binding Protein-MBP tagged proteins), or Spycatcher/Snoopcatcher^27^ (to bind to SpyTag/SnoopTag-tagged proteins).

An additional major advantage of our method is that it ensures that micropatterned proteins maintain their activity, which was best exemplified by our successful micropatterning of active molecular motors (Fig. 3). This is especially important for sequential micropatterning using LIMAP, as the ROS generated by UV light to micropattern the second protein can potentially be harmful to the first micropatterned protein. This improvement in activity is achieved by first micropatterning Fibrinogen anchors that specifically binds to purification tags, followed by addition of the tagged proteins of interest. As the proteins of interest are added after the actual patterning process, they can be added in the optimal buffer for their stability/activity. Conversely, the Fibrinogen anchor is micropatterned in its optimal buffer for patterning efficiency, thereby removing the need to compromise between the two. It must be emphasized that our technology also maximises activity by mitigating preferential orientation effects as it ensures that binding to the pattern occurs at the level of the protein tag, usually added to flexible N- or C-terminal regions, rather than on the “stickiest” face of the protein, which could be (next to) the active site. In particular, this allowed us to micropattern molecular motors specifically via their C-terminal tail away from their N-terminal motor domain, thereby maintaining their activity (Fig. 3). We envision that these key assets will provide major advantages for *in vivo* studies, to micropattern active ligands in the right orientation to bind to membrane receptors, but also for *in vitro* studies, for multiplexed micropatterning of various molecular motors to pave the way towards the *in vitro* reconstitution of complex cytoskeleton landscapes, akin to those found in cells.

In addition to ensuring protein activity, our method also minimizes cross-adsorption between successive patterns as it simplifies pattern quenching. In other words, not only are the proteins active, but also they are at the right place (see figure 2). Indeed, because it is virtually always the same protein that is patterned, this simplifies the determination of the quenching conditions for each specific experiment. Coupled to the constant homogeneity between patterns brought by our method, we envision that this property will be a key advantage for multi-protein applications where control over the relative patterning density between proteins is key, for example in testing the cellular response to multiple gradients of signalling molecules.

The combination of all the advantages offered by our technology enabled us to achieve subcellular micropatterning of proteins, whereby the position of proteins can be imprinted from the outside via a micropattern (Fig. 5, 6). This was not possible with direct patterning of ligands (Fig. S7), demonstrating the importance of the fibrinogen toolbox described here. While beyond the scope of this paper, we envision that this technology will allow researchers to untangle the interplay between mechanical forces (provided by the distribution of adhesion molecules) and signalling (provided by the distribution of signalling receptors). Conversely, by trapping endogenous receptor-ligand complexes into the TIRF field while blocking their endocytosis, this assay could help the characterization of the sequence of recruitment of proteins at endocytic sites, which could be then structurally characterized thanks to the recent combination of CryoEM and micropatterning^9,10^. Furthermore, the reconstitution of signalling platforms of defined receptor-ligand content at a known density is poised to help deciphering the combinatorial interactions between signalling pathways, such as the cross-talks between EGFR and Her2 which has been proposed to underlie resistance to anticancer drugs targeting these receptors^28^. Last, the orthogonal transmembrane segment we established (Fig. 5) could be used to relocalise other proteins to the cell cortex in a controllable fashion via a micropattern, an assay reminiscent to what we previously achieved *in vivo* using polarity markers as an anchoring platform^29^. This could for instance be used to generate symmetry breaking of the actin cortex by asymmetric targeting of cytoskeleton regulators (by fusing cytoskeleton regulators to the transmembrane construct). In conclusion, we hope that the advantages of the Fibrinogen anchors described in this paper will be a stepping stone towards having micropatterning experiments only limited by the imagination of the researcher rather than by intrinsic limitations of this technology.

## Methods

### Synthesis of the photosensitizer (4-benzoylbenzyl)trimethylammonium bromide: BBTB)

All chemical reagents were purchased with the highest purity available. Briefly, in a 100 mL round bottom flask, 2.1 mL trimethylamine solution 4.2M in ethanol (1.2 eq, 8.8 mmol, Aldrich) was added to 2.0 g of 4-(bromomethyl)benzophenone (1 eq, 7.3 mmol, Aldrich) resuspended in 40 mL Acetonitrile. The reaction was placed under reflux for 2h in a nitrogen atmosphere. Once the reaction was completed, the solvent was evaporated, and the white solid obtained was dried under vacuum (2.35 g, 97% yield).

HPLC > 95% pure

^1^H NMR (400 MHz, MeOD) δ: 7.93 (m, 2H), 7.84 (m, 2H), 7.77 (m, 2H), 7.70 (m, 1H), 7.58 (m, 2H), 4.71 (s, 2H), 3.22 (s, 9H)

^13^C NMR (400 MHz, MeOD) δ: 139.5, 136.8, 132.8, 132.8, 131.7, 130.0, 129.7, 128.3, 68.3, 52.0 MS (ESI) [m-Br]^+^/z Calculated for C_17_H_20_NO: 254.1, found: 254.1

### Plasmids

The GBP3xCys consists of a nanobody against GFP^21^, referred to as GBP, with three cysteines added far from the active site to minimize loss of activity upon functionalization through these cysteines^25^. GBP3xCys was amplified by PCR from the P434_H14-Sp-brNEDD8-Cys-anti-GFP nanobody (a kind gift from Tino Pleiner) and cloned into a modified pRSET vector (a kind gift from Mark Allen) tagging N-terminally the ORF with a (His)_6_ tag, followed by the first 93 amino acids of the dihydrolipoyl acetyltransferase acetyltransferase component of pyruvate dehydrogenase complex from *Bacillus stearothermophilus* to enhance solubility (as previously described^30^), and a TEV cleavage site (referred to as His-SS-TEV). EGFP was PCR-amplified from pEGFP-C2 (Clontech) and similarly cloned into the pRSET His-SS-TEV vector. pWC2^23^ encoding the motor domain of *Drosophila* kinesin 1 fused to BCCP (noted Kin1-biotin) was a gift from Jeff Gelles (Addgene plasmid # 15960; http://n2t.net/addgene:15960; RRID:Addgene_15960).

For the purification of Streptavidin-GFP-GFP, two GFPs were cloned into the pRSET His-SS-TEV-Streptavidin, which permits solubilisation of natively-purified Streptavidin^30^. The GFP sequences were codon optimised to prevent repetitive sequences.

The transmembrane nanobody construct (Fig. 5) comprises an N-terminal signal peptide from the *Drosophila* Echinoid protein, followed by (His)_6_-PC tandem affinity tags, the GBP nanobody against GFP^21^, a TEV cleavage site, the transmembrane domain from the *Drosophila* Echinoid protein, the VSV-G export sequence and the mScarlet protein^31^. The protein expressed by this construct thus consists of an extracellular anti-GFP nanobody linked to an intracellular mScarlet by a transmembrane domain (named GBP-TM-mScarlet in the main text for simplicity). This custom construct was synthesized by IDT and cloned into a modified pCDNA5/FRT/V5-His vector, as previously described^32^ for homologous recombination into the FRT site.

### Proteins

Unless stated otherwise, all protein purification steps were performed at 4°C. Protein concentration was determined either by absorbance at 280nm, or by densitometry on coomassie-stained SDS page gel against a BSA ladder.

GBP3xCys, Streptavidin-GFP-GFP and GFP were expressed in *E. coli* BL21 Rosetta 2 (Stratagene) using the plasmids described above by induction at OD600=0.8 with 1 mM IPTG in 2X YT (Streptavidin-GFP-GFP) or TB (GFP, GBP3xCys) medium at 20°C. Bacteria were lysed by sonication in lysis buffer (20 mM Hepes, 150 mM KCl, 1% Triton X-100, 5% Glycerol, 20 mM imidazole, 5 mM MgCl_2_, 0.5 mM DTT, pH 7.6) enriched with protease inhibitors (Roche Mini) and 1 mg/ml lysozyme (Sigma) and 10 μg/ml DNAse I (Roche). After clarification (16,000 rpm, Beckman JA 25.5, 30 min, 4°C), lysate was incubated with Ni-NTA resin (Qiagen) for 2 h at 4°C and washed extensively in 20 mM Hepes, 150 mM KCl, 5% glycerol, 15 mM imidazole, 0.5 mM DTT, pH 7.6). Protein was eluted in 20 mM Hepes, 150 mM KCl, 5% glycerol, 250 mM imidazole, 0.5 mM DTT, pH 7.6. Bradford-positive elution fractions were pooled and TEV-cleaved overnight by adding 1:50 (vol:vol) of 2 mg/ml (His)_6_-TEV protease and 1 mM/0.5 mM final DTT/EDTA. The solution was then dialysed twice against 20 mM Hepes, 150 mM KCl, 15 mM Imidazole, 0.5 mM DTT, pH 7.7, and TEV protease and His-SS-TEV were removed using Ni-NTA resin. Tag-free GBP3xCys was then concentrated down to 10 mg/ml, flash frozen in liquid N_2_ and kept at −80°C. GFP and Streptavidin-GFP-GFP were similarly purified, except that DTT was omitted in the lysis and storage buffers, and that final concentration was 4.22 mg/ml (GFP) and 0.75 mg/ml (Streptavidin-GFP-GFP).

BSA-LC-LC-Biotin-Alexa647 (BSA-Biotin-Alexa647) was generated by reacting 30 μM Bovine Serum Albumin (BSA, Fisher, BP1605, dissolved in 0.1 M sodium bicarbonate, pH 8.3) simultaneously with a 5-fold molar excess of Alexa Fluor 647 NHS Ester (ThermoFisher, A20006) and a 5-fold molar excess of EZ-Link NHS LC-LC-Biotin (ThermoFisher, 21343). Both NHS-reactive chemicals were resuspended immediately prior to the reaction in a new vial of anhydrous dimethylsulfoxide (DMSO, Invitrogen, D12345). The reaction was rocked for one hour at room temperature, before removal of unreacted Alexa647 and biotin on a Zeba Spin column (ThermoFisher, 89891) equilibrated in 0.1 M sodium bicarbonate, pH 8.3. The degree of Alexa647 labelling was 1.48 mol of dye per mol of BSA, and the resulting BSA-Biotin-Alexa647 visually bound to agarose beads conjugated to streptavidin (Pierce, 20359).

To generate streptavidin-Alexa647, streptavidin (1mg/ml in PBS + 60 mM sodium bicarbonate, pH 8.0) was reacted with a 6-fold molar excess of Alexa Fluor 647 NHS Ester (ThermoFisher, A20006). Excess dye was removed by Zeba Spin column equilibrated in 20 mM Hepes, 150 mM KCl, 5% glycerol pH 7.6.

Kin1-biotin was expressed using the plasmid pWC2 by induction at OD600=0.8 with 1 mM IPTG in 2X YT medium at 20°C. Bacteria were lysed by sonication in lysis buffer (20 mM Hepes, 150 mM KCl, 1% Triton X-100, 5% Glycerol, 0.1 mM ATP, 10 mM imidazole, 10 mM MgCl_2_, 1 mM DTT, pH 7.6) enriched with protease inhibitors (Roche Mini) and 0.7 mg/ml lysozyme (Sigma) and 10 μg/ml DNAse I (Roche). After clarification (14,000 rpm, Beckman JA 25.5, 30 min, 4°C), lysate was incubated with Ni-NTA resin (Qiagen) for 2 h at 4°C and washed extensively in wash buffer (20 mM Hepes, 150 mM KCl, 5% Glycerol, 0.1 mM ATP, 10 mM imidazole, 2 mM MgCl_2_, 1 mM DTT, pH 7.6), followed by wash buffer enriched with 2 mM ATP and a final wash in wash buffer. Protein was eluted in 20 mM Hepes, 150 mM KCl, 5% Glycerol, 0.1 mM ATP, 300 mM imidazole, 2 mM MgCl_2_, 1 mM DTT, pH 7.6. Bradford-positive elution fractions were pooled and diluted 1:11 (vol eluate:vol QA) in buffer QA (20 mM Hepes, 20 mM KCl, 1 mM MgCl_2_, 0.05 mM ATP, 1 mM DTT, pH 7.6) before loading onto an MonoQ 5/50 GL anion exchange column. Protein was then eluted in a 0.05-1M gradient of KCl. Positive fractions were pooled, dialysed against storage buffer (20 mM Hepes pH 7.7, 150 mM KCl, 1 mM MgCl_2_, 0.05 mM ATP, 1 mM DTT, 20% (w/v) sucrose, pH 7.6), flash frozen in liquid N_2_ and kept at −80°C (final concentration 2.2 mg/ml).

NeutrAvidin, NeutrAvidin DyLight-550, and Fibrinogen-Alexa546 were purchased from Thermofisher. Unlabelled Fibrinogen was purchased from MP Biomedicals (#08820224).

Tubulin, HiLyte-647 tubulin and biotinylated tubulin were all purchased from Cytoskeleton. GMPCPP-stabilised microtubules were polymerised by resuspending 1 μl aliquots of HiLyte-647 labelled tubulin mixes (60% unlabelled tubulin, 20% HiLyte-647 tubulin, 20% biotinylated tubulin, Cytoskeleton, Inc. T240, TL670M, T333, respectively) in 50 μl of warm (80 mM PIPES, 2 mM MgCl2, 0.5 mM EGTA, 0.6 mM GMPCPP, pH 6.9), before incubation for 30 minutes at 37°C. Microtubules were pelleted for 8 minutes at 15,871 x *g* and resuspended in 50 μl of (80 mM PIPES, 2 mM MgCl2, 0.5 mM EGTA, 0.6 mM GMPCPP, pH 6.9).

Delta-like ligand 4 (DLL4), comprising a fragment of the human Delta ectodomain (1-405) with a C-terminal GS-SpyTag-His_6_ sequence, was purified from culture medium of transiently transfected Expi293F cells (Thermo Fisher) by metal affinity chromatography. Protein was further purified by size exclusion chromatography on a Superdex 200 column in 50 mM Tris pH 8.0, 150 mM NaCl and 5% glycerol. DLL4 (69 μM) was subsequently buffer-exchanged into 0.1 M sodium bicarbonate, pH 8.3 using a Zeba spin column, and labelled with a 3-fold molar excess of EZ-Link NHS LC LC Biotin (ThermoFisher) for 1h at RT. Unreacted biotin was then removed by Zeba Spin column equilibrated in 0.1 M sodium bicarbonate, pH 8.3.

### Fibrinogen fusions

#### 3 step synthesis of Fibrinogen-NeutrAvidin, Fibrinogen GFP and Fibrinogen ConA

Fibrinogen powder was resuspended at 4 mg/ml (~11.8 μM) in Fibrinogen buffer (0.1 M sodium bicarbonate, 0.5 mM EDTA, pH 8.3), before addition of a 25-fold molar excess of Traut’s Reagent (2-iminothiolane, Pierce). The solution was gently rocked for 45 min at RT. 1 mM DTT final was then added and the solution was further rocked for 1h at RT. Excess DTT/ Traut’s reagent was then removed using a Zeba Spin column equilibrated in Fibrinogen buffer. Concomitantly, the buffer of purified NeutrAvidin (80 uM) was exchanged for fresh Fibrinogen buffer using a Zeba Spin column, then a 6-fold molar excess of Maleimide-PEG8-succinimidyl ester (Sigma 746207) was added. After rocking for 1h at RT, the excess Maleimide-PEG8-succinimidyl ester was removed on a Zeba Spin column equilibrated in Fibrinogen buffer. NeutrAvidin-Maleimide was then added to Fibrinogen-Thiol in a 4-fold NeutrAvidin-maleimide::Fibrinogen molar ratio and rocked overnight at 4°C. 50 mM final free cysteine was then added and incubated for 30 min at RT to quench the remaining unreacted maleimide functions. The solution was then brought to 25% final ammonium sulfate by adding one-third of the volume of saturated ammonium sulfate to specifically precipitate the Fibrinogen-NeutrAvidin. After 30 min rocking at RT, the solution was centrifuged at 16,873 x *g* for 10 min. The pellet was resuspended in the initial volume of Fibrinogen buffer by gentle rocking for 1h at 4°C, followed by a second precipitation in 25% final ammonium sulfate as above. After centrifugation at 16,873 x *g* for 10 min, the pellet was resuspended in one quarter of the initial volume of Fibrinogen buffer, ultracentrifuged to remove aggregates (100,000 x *g*, 5 min at 4°C) then flash frozen in liquid N_2_ and kept at −80°C.

Fibrinogen-GFP was similarly obtained by replacing NeutrAvidin by purified GFP and in this case, we used a 15-fold molar excess of GFP-Maleimide over Fibrinogen-Thiol. The absorbance at 488nm of GFP allowed us to evaluate the degree of labelling to 0.51 mol of GFP per mol of Fibrinogen in these conditions.

Fibrinogen-ConA was similarly obtained by replacing NeutrAvidin by purified ConA (Sigma).

#### 2 step synthesis of Fibrinogen-GBP

Fibrinogen powder was resuspended at 4mg/ml (~11.8 μM) in Fibrinogen buffer (0.1 M sodium bicarbonate, 0.5 mM EDTA, pH 8.3), then a 25-fold molar excess of Maleimide-PEG8-succinimidyl ester was added. After rocking for 1h at RT, the excess Maleimide-PEG8-succinimidyl ester was removed on a Zeba Spin column equilibrated in Fibrinogen buffer. Concomitantly, the buffer of purified GBP3xCys was exchanged for fresh Fibrinogen buffer using a Zeba Spin column, which also removed the DTT used in the purification of the GBP3xCys to keep cysteines reduced. GBP3xCys (10 mg/ml) was then added to Fibrinogen-maleimide in a 5-fold GBP::Fibrinogen-maleimide molar ratio and rocked overnight at 4°C. 50 mM (final) free cysteine was then added and incubated for 30 min at RT to quench the remaining unreacted maleimide functions. Then the solution was brought to 25% final ammonium sulfate as above to specifically precipitate the Fibrinogen-GBP. After 30 min rocking at RT, the solution was centrifuged at 16,873 x *g* for 10 min and the pellet was resuspended in the initial volume of Fibrinogen, followed by a second precipitation in 25% final ammonium sulfate. After centrifugation at 16,873 x *g* for 10 min, the pellet was resuspended in one quarter of the initial volume of Fibrinogen buffer, ultracentrifuged to remove aggregates (100,000 x g, 5 min at 4°C) then flash frozen in liquid N_2_ and kept at −80°C.

#### One step synthesis of Fibrinogen-biotin, Fibrinogen-Alexa647, Fibrinogen-Biotin-ATTO490LS and Fibrinogen-Biotin-Alexa647

Fibrinogen-biotin was obtained by mixing Fibrinogen (4 mg/ml in Fibrinogen buffer: 0.1 M sodium bicarbonate, 0.5 mM EDTA, pH 8.3) with a 20-fold molar excess of EZ-Link NHS LC LC Biotin (Pierce) for 1h at RT. Unreacted biotin was then removed by dialysis against Fibrinogen buffer (2×2L).

Fibrinogen-Alexa647 was obtained by reacting Fibrinogen (4 mg/ml in Fibrinogen buffer) with a 10-fold molar excess of NHS Alexa 647 (ThermoFisher, A20006) for 1h at RT. Unreacted dye was then removed by Zeba Spin column equilibrated in Fibrinogen buffer.

Fibrinogen-biotin-ATTO490LS and Fibrinogen-biotin-Alexa647 were generated by mixing Fibrinogen (4mg/ml in Fibrinogen buffer) with a 3-fold molar excess of ATTO490LS NHS ester (ATTO-TEC) or Alexa647 NHS ester (ThermoFisher, A20006) for 15 minutes at RT. The Fibrinogen-ATTO490LS and Fibrinogen-Alexa647 were subsequently reacted with a 50-fold molar excess of EZ-Link NHS LC LC Biotin for 1h at RT. Unreacted biotin and fluorescent dye were then removed by Zeba Spin column equilibrated in Fibrinogen buffer.

### Cells

Drosophila S2 cells (UCSF, mycoplasm-free judged by DAPI staining) were grown in Schneider Medium (Gibco) supplemented with 10% heat-inactivated Foetal Bovine Serum (FBS). Flp-In NIH/3T3 cells (Invitrogen) were cultured in DMEM (Gibco) supplemented with 10% Donor Bovine Serum (Gibco) and Pen/Strep 100 units/ml at 37°C with 5% CO_2_. HeLa cells (ATCC, CCL-2) were cultured in DMEM supplemented with 10% Foetal Bovine Serum (FBS, Gibco) and Pen/Strep 100 units/ml at 37°C with 5% CO_2_. U2OS cells (ATCC, HTB-96) stably expressing inducible FLAG-Notch1-EGFP chimeric receptors^33^ were maintained in DMEM supplemented with 10% FBS, Pen/Strep 100 units/ml, 50 μg/ml hygromycin B (Thermo) and 15 μg/ml blasticidin (Invitrogen) at 37°C with 5% CO_2_. Prior to use in experiments, U2OS cells were induced with 2 μg/ml doxycycline for 24 hours. Flp-In NIH/3T3 were transfected with a modified pCDNA5 vector containing GBP-TM-mScarlet with Lipofectamine 2000 (Invitrogen). Stable transfectants were obtained according to the manufacturer’s instructions by homologous recombination at the FRT were selected using 100 μg/ml Hygromycin B Gold (Invivogen). All live imaging was performed in Leibovitz’s L-15 medium (Gibco, 11415064) supplemented with 10% Donor Bovine Serum and HEPES (Gibco, <u>1563080</u>, 20 mM).

### Immunofluorescence

Cells were fixed 4 % paraformaldehyde in PBS for 20 min, permeabilized with 0.1 % Triton X-100 in PBS for 5 min, then washed in PBS, then in PBS supplemented with 1% BSA and 1% rabbit IgG blocking reagent (ThermoFisher) for 5 min, then in PBS. Anti-Clathrin (Abcam, ab21679) and anti-EGF Receptor (Cell Signaling, D38B1) were labelled with Alexa Fluor 488- and Alexa Fluor 647-Zenon Rabbit IgG labelling kits respectively (ThermoFisher), according to the manufacturer’s instructions, and cells were then incubated with both antibodies (both diluted 1:100 in PBS-1% BSA) for 20 minutes. After washing thrice in PBS, imaging was performed in PBS instead of mounting medium to avoid squashing the cells (and potentially biasing the Clathrin or EGFR / pattern colocalization).

### Microscopy

Imaging was performed on a custom TIRF/spinning disk confocal microscope composed of a Nikon Ti stand equipped with perfect focus and a 100X NA 1.45 Plan Apochromat lambda objective (or alternatively, a 60X NA 1.49 Apochromat TIRF or a 10X NA 0.3 Plan Fluor). The confocal imaging arm is composed of a Yokogawa CSU-X1 spinning disk head and a Photometrics 95B back-illuminated sCMOS camera operating in global shutter mode and synchronized with the spinning disk rotation. Conversely, the TIRF imaging arm is composed of an azimuthal TIRF illuminator (iLas2, GATACA systems) modified to have an extended field of view (Cairn) to match the full field of view of the camera. Images are recorded with a Photometrics Prime 95B back-illuminated sCMOS camera run in pseudo-global shutter mode and synchronised with the azimuthal illumination. Excitation is performed using 488 (150mW OBIS LX), 561 (100mW OBIS LS) and 637 nm (140mW OBIS LX) lasers fibered within a Cairn laser launch. To minimise bleed through, single band emission filters are used (Chroma 525/50 for GFP, 595/50 for Alexa546 and ET655lp for Alexa647/ATTO647N/ATTO490LS/SiR-Tubulin) and acquisition of each channel is performed sequentially using a fast filter wheel (Cairn Optospin) in each imaging arm. Filter wheels also contain a quad-band filter (Chroma ZET405/488/561/640m) to allow imaging of the reflection of the DMD illumination arm at the glass/water interface (see below). To enable fast acquisition, the entire setup is synchronized at the hardware level using an FPGA stand-alone card (National Instrument sbRIO 9637) running custom code. TIRF angle was set independently for all channels so that the depth of the TIRF field was identical for all channels. Sample temperature was maintained at 25°C using a heating enclosure (MicroscopeHeaters.com, Brighton, UK). Acquisition was controlled by Metamorph software.

### Optical design of a low-cost UV-LED DMD illuminator

Our optical design combines DMD-UV-illumination with TIRF illumination onto the Nikon Ti setup described above (see Fig. S1). Briefly, a 385 nm high power UV LED light source (M385LP1, Thorlabs) is collimated using an AR coated aspheric lens (ACL2520U-A, Thorlabs). The collimated UV beam is then directed towards a Digital Micromirror Device (DMD, DLPLCR6500EVM, Texas Instrument) at an 24° angle of incidence (corresponding to twice the tilting angle of the DMD mirrors). The Image of the DMD chip is then relayed onto the conjugate of the sample plane at the backport of the microscope through a 4f imaging system (f1 = f2 = 125 mm UV Fused Silica Bi-Convex Lenses, AR-Coated, LB4913-UV, Thorlabs). This intermediate image is then relayed onto the sample plane by a tube lens (125 mm, UV rated, Edmond optics) and the objective (100X Plan Apochromat lambda NA 1.45, Plan Apochromat 60X NA 1.4, or 20X Plan Apochromat Lambda VC NA 0.75). To combine DMD-UV illumination with TIRF-illumination, an ultraflat dichroic (T470lpxr, Chroma) is placed after f2 within a custom backport assembly (Cairn). When in DMD UV illumination mode, the microscope filter cube turret contains a 473nm dichroic (Di03-R473 Semrock) while it contains an ultra-flat quad band dichroic/clean-up filter (Chroma TRF89901-EM) when in TIRF illumination mode. The 473nm dichroic is compatible with simultaneous spinning disk/ DMD illumination. We note that care must be taken with the adjustment of the collimating lens of the LEDs to find the best compromise between illumination intensity and flatness of the illumination profile. If necessary, a flatfield correction can be applied on the patterns to be displayed to account for any field inhomogeneities, and if extremely sharp patterns are required, an iris can be put in the Fourier plane between f1 and f2, but we found that this was not required for most applications.

To offer a second illumination wavelength for 450nm optogenetic stimulation using the same DMD chip, our design also contains a second collimated LED (450nm M450LP1 Thorlabs in our case) at the symmetric −24° angle. Therefore, any pattern can be displayed using the 450nm light source by simply inverting the pattern before displaying it on the DMD. This could be replaced by any other light source to bring epifluorescence imaging in order to facilitate multiprotein pattern alignment on a setup not equipped with TIRF.

LED intensity is controlled using a custom LED driver providing the maximum 1.7A tolerated by the LED (respectively 2A for 450nm LED). Control over LED intensity and on/off state is operated using a Digital/Analog card (National Instrument USB-6001 or Arduino UNO equipped with a custom shield providing a Texas Instrument TLV5618 DAC). Communication to the DMD from the imaging software is performed using the DMD connect library developed by Klaus Hueck^34^ (https://github.com/deichrenner/DMDConnect). Control of all parts was integrated into Metamorph and/or Micromanager using custom scripts to calibrate the DMD with respect to the camera, display user-defined UV patterns and facilitate pattern alignment for multiprotein patterning.

To keep the cost low and to enable researchers with limited access to mechanical workshops to make their own module, our design relies on a commercially available cage system to set the +24° angle and a custom mount for the DMD board, which can be machined or 3D printed. This greatly facilitates alignment of the setup, as this essentially locks all pieces into the correct angle during assembly, which is critical for alignment of DMD setups^16^. All codes, and CAD files for this setup are available freely upon request for non-commercial purposes.

Importantly, efficient and crisp patterning requires that the microscope is focused on the PLL-PEG (or PEG-Silane)/Glass interface. While hardware autofocus systems help in finding this interface, they do not always work perfectly as there is usually a correction offset to add, which may vary from sample to sample. To ensure that we always focus the instrument at the right place, we use the fact that because patterning is performed in aqueous buffer, there is a glass/liquid interface at the PLL-PEG layer, which reflects the UV patterning light. We thus chose our filter/dichroic sets to allow some of this reflected light to be imaged onto the camera, which allows us to easily find the optimal sample plane in the absence of anything fluorescent in the chamber. Once this plane has been found, we activate the hardware autofocus system to ensure that this plane is kept during patterning process.

### Protein Micropatterning

For micropatterning on PLL-PEG-passivated coverslips in an open configuration (all patterns except Fig. S4), clean-room grade coverslips (Nexterion, custom 25 x 75 mm size) were surface activated under pure oxygen in a plasma cleaner (PlasmaPrep2, GaLa instruments) and then laid on top of a 200 μl drop of filtered PLL(20)-g[3.5]-PEG(2) PLL-PEG (PLL-PEG, SuSoS) (100 μg/ml in 10 mM Hepes, pH 7.6) for 1 hour, in a humid chamber. Coverslips were then washed extensively in filtered MilliQ water, and dried under a flow of dry nitrogen gas. Coverslips were then mounted in a sticky slide Ibidi 8-well chamber (Ibidi, 80828) and pressed under a ~4 kg weight overnight to stick the chamber to the glass. Then, per well of the 8-well chamber, 200 μl of BBTB was added (50 mM in 0.1 M sodium bicarbonate pH 8.3) and exposed to patterned UV light using the DMD-UV arm of the micropatterning microscope (30-90 s exposure) after focussing on the glass/buffer interface. BBTB was subsequently removed by repeated dilution (twelve washes with 600 μl of 0.1 M sodium bicarbonate, pH 8.3), before addition of 200 μl of Fibrinogen anchor (100 μg/ml in 0.1 M sodium bicarbonate, pH 8.3, for a final concentration of 50 μg/ml). The Fibrinogen anchor was allowed to adsorb to the surface for 5 minutes, before washing by repeated dilution (12 washes). The surface was then quenched for 5 minutes with of 0.5 mg/ml PLL-PEG (in 10 mM Hepes pH 7.6). PLL-PEG was then removed by similar repeated dilution, so that the surface never dried. The protein of interest was then added it its buffer of choice (here 0.1 M sodium bicarbonate pH 8.3 for GFP, biotin-BSA and NeutrAvidin) at 5 μg/ml and allowed to bind to the fibrinogen anchor for 5 min before extensive washes and imaging. Alternatively, when proteins were directly patterned, the Fibrinogen anchor in the above protocol was replaced by the protein of interest at a concentration of 50 μg/ml in 0.1 M sodium bicarbonate pH 8.3 (except for fluorescent EGF, where biotinylated EGF complexed to streptavidin-Alexa555 (Invitrogen, E35350) was pattered at 1 μg/ml in 0.1 M sodium bicarbonate, pH 8.3). After a 5 min incubation, the surface was quenched and washed as above.

For dual patterning experiments, presented in Fig. 2, on PLL-PEG-passivated coverslips, the protocol above was performed, except that immediately after the PEG quenching step, BBTB was added to the chamber (50 mM in 0.1 M sodium bicarbonate pH 8.3) and the whole process was repeated for a second Fibrinogen anchor. The two proteins of interest were then added together onto the pattern for 5 min (5 μg/ml each in 0.1 M sodium bicarbonate, pH 8.3), before extensive washes and imaging. All Fibrinogen anchors that are not fluorescent, were doped with 5 μg/ml Fibrinogen-Alexa546 to image their respective pattern and thereby align the next pattern. For controls where one Fibrinogen anchor was omitted, it was replaced with Fibrinogen Alexa546.

For dual patterning experiments where receptors were clustered to micropatterns (Fig. 5, 6, S7), the above protocol was performed, with the small central region patterned first (50 μg/ml GFP, Fibrinogen-GFP, Fibrinogen-Biotin-ATTO490LS or Fibrinogen-Biotin-Alexa647), before PLL-PEG quenching and patterning of the second, outer region (50 μg/ml Fibronectin + 10 μg/ml Fibrinogen-Alexa546 or Fibrinogen-Alexa647). For relocalisation of the GBP-TM-mScarlet receptor, patterns (with a central Fibrinogen-Biotin-ATTO490LS centre) were incubated (or not) with Streptavidin-GFP-GFP (10 μg/ml) and fixed with 0.5 mM dithiobis(succinimidyl propionate)) (DSP) for 20 minutes. After extensive washing, NIH/3T3 cells were added (20,000 cells/well) in serum-free DMEM for 30 minutes, before washing into medium containing serum, and imaging 30 minutes later. Alternatively, for live imaging of GBP-TM-mScarlet recruitment (Figure 5), 20,000 cells were added per well, in L15 + 20 mM Hepes, and imaged as they landed and spread on micropatterns. For relocalisation of EGFR and GFP-Notch1, double patterns were incubated with NeutrAvidin (25 μg/ml) for 5 minutes, before extensive washing and incubation for 5 minutes with biotinylated-EGF (1 μg/ml) or biotinylated-DLL4 (0.5 μM) respectively. Chambers were then extensively washed. HeLa cells were serum starved for 24 hours before addition, in serum-free medium, to EGF-micropatterns (20,000 cells/well). Cells were left to spread for 40 minutes prior to fixation. U2OS GFP-Notch1 cells, induced for 24 to express GFP-Notch1, were added to DLL4-micropatterns in L15 + 20 mM Hepes (20,000 cells/well) and imaged as they landed and spread on micropatterns.

For patterning of Kin1-biotin using the Fibrinogen-biotin/NeutrAvidin “sandwich” (Fig. 3), PLL-PEG-passivated coverslips assembled into 8-well chambers were exposed for 90s, before the addition of Fibrinogen-Biotin-ATTO490LS (50 μg/ml in 0.1 M sodium bicarbonate, pH 8.3) as above. Patterns were then quenched with 0.5 mg/ml PLL-PEG for 5 minutes before addition of NeutrAvidin (10 μg/ml in 0.1 M sodium bicarbonate pH 8.3 for 5 minutes), before washing and incubation with 5 μg/ml purified Kin1-biotin in (80 mM PIPES, 1 mM MgCl2, 5 mM ATP, pH 8.9) for 5 minutes. Alternatively, for direct patterning of Kin1-biotin, patterns were exposed to patterned UV light in the presence of BBTB for 90 s as above before addition of 5 μg/ml Kin1-biotin in (80 mM PIPES, 1 mM MgCl2, 5 mM ATP, pH 8.9). Micropatterns were then quenched with 0.5 mg/ml PLL-PEG for 5 minutes. 4 μl of GMPCPP MT seeds (see above) were then added to each well, in 200 μl of ATP-regenerating buffer (80 mM PIPES, 0.1 mg/ml κ-casein, 40 μM DTT, 64 μM glucose, 160 μg/ml glucose oxidase, 20 μg/ml catalase, 0.1% methylcellulose, 1 mM ATP, pH 6.9), and imaged by TIRF microscopy. For direct patterning of Kin1-biotin, visualisation of patterns was performed post imaging, by addition of 1 μM streptavidin-Alexa647.

For micropatterning of ConA to bind to S2 cells (Fig. 4), surface-activated coverslips were passivated with 0.1 mg/ml PLL-PEG before being patterned using a quartz-chromium photomask and a deep UV light source for 5 minutes as described previously^11^. Exposed coverslips were then incubated with 200 μl Fibrinogen alone (50 μg/ml Fibrinogen, 5 μg/ml Fibrinogen-Alexa546), ConA alone (50 μg/ml ConA, 5 μg/ml Rhodamine-ConA) or Fibrinogen-ConA (50 μg/ml Fibrinogen-ConA, 5 μg/ml Fibrinogen-Alexa546) all in 0.1 M sodium bicarbonate pH 8.3 for 1 hour. Coverslips were washed in 0.1 M sodium bicarbonate pH8.3 and assembled into an 8-well chamber. 100,000 S2 cells in low (1%) serum media, were then added to each well for 1 hour, before incubation with SiR-Tubulin^35^ (1 μM, Spirochrome) for 30 minutes to label cells. Chambers were washed with Schneider medium + 10% heat-inactivated FBS and imaged by confocal spinning disc microscopy.

For micropatterning on PEG-silane-passivated coverslips in a flow cell configuration (Fig. S4), clean-room grade coverslips (Nexterion, 75 x 25 mm) were surface activated under pure oxygen in a plasma cleaner and then incubated with PEG-silane (30 kDa, PSB-2014, creative PEG works) at 1 mg/ml in ethanol 96% /0.1% of HCl, overnight at room temperature with gentle agitation. Standard 22×22 mm coverslips were similarly passivated with PEG-silane, omitting the plasma-cleaning step. Slides and coverslips were then successively washed in 96% ethanol and ultrapure water, before drying under nitrogen gas. The coverslip and slide were subsequently assembled into an array of six flow cells (~15μL each) using double sided tape (Adhesive Research AR-90880 cut with a Graphtec CE6000 cutting plotter). The flow cell chamber was then filled with BBTB (50 mM in 0.1 M sodium bicarbonate pH 8.3) and exposed to UV light on the DMD-UV arm of the micropatterning microscope (3s exposure) after focussing on the glass/buffer interface. The flow cell was subsequently washed with three flow cell volumes of carbonate buffer, and Fibrinogen-biotin was then injected at 20 μg/ml in 0.1 M sodium bicarbonate pH 8.3. After a two-minute incubation, the chamber was washed with three flow cell volumes of carbonate buffer, and NeutrAvidin (or NeutrAvidin-Dylight-550) was injected at 50 μg/ml in 0.1 M sodium bicarbonate pH 8.3. The chamber was then washed with Hepes buffer (10 mM Hepes, pH 7.6) and quenched with PLL-PEG (0.2 mg/ml in 10 mM Hepes, pH 7.6) for two minutes. The chamber was then washed with Hepes buffer and BSA-Biotin-Alexa647 was added (10 μg/ml in 10 mM Hepes, pH 7.6) for one minute, before extensive washing with Hepes Buffer before imaging. For controls where NeutrAvidin was directly patterned, Fibrinogen-biotin was replaced by NeutrAvidin/NeutrAvidin-DyLyte550 (50 μg/ml in carbonate buffer), and the chamber was directly washed with Hepes buffer, quenched with PLL-PEG and incubated with BSA-Biotin-Alexa647 as above.

In general, we found that buffer composition (see Fig. S2) and avoiding drying of the surface (see Fig. S5) were fundamental for ensuring reproducibility, selectivity and homogeneity of micropatterning. Note that while the patterning buffer is important, the use of Fibrinogen anchors allows one to bind the proteins of interest to the pattern in virtually any buffer, as this binding step occurs after the patterning process.

### Image processing

Images were processed using Fiji^36^. Figures were assembled in Adobe Illustrator 2019.

Patterning selectivity and homogeneity were computed as follows:

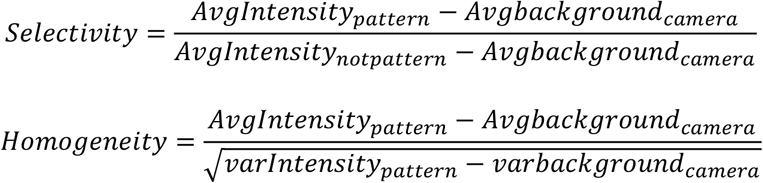

With *AvgIntensity_pattern_* representing the average fluorescence Intensity of the fluorescent protein onto the pattern and its associated variance *varIntensity_pattern_, AvgIntensity_notpattern_* representing the average fluorescence Intensity of the fluorescent protein onto a non-patterned region adjacent to the pattern, *Avgbackground_camera_* representing the average background Intensity of the camera and its associated variance *varbackground_camera_*. *AvgIntensity_pattern_, AvgIntensity_notpattern_* and *varIntensity_pattern_* were measured in a Region of Interest of identical size (and as large as possible to provide good estimates). *Avgbackground_camera_* and *varbackground_camera_* were obtained by measuring the signal in the dark upon screwing a lid onto the camera.

Similarly, we measured a proxy of the amount of protein specifically being deposited onto the patterned area as:

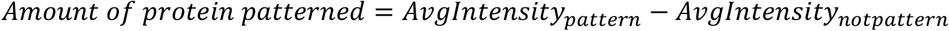

Note that while the *Selectivity* is a normalized value and can be compared between fluorophores, the *Amount of protein patterned* is only valid when comparing the same fluorophore as if will also depend on the photophysics of the dye and the sensitivity of the instrument to the dye.

Similarly, the enrichment of mScarlet relocalisation (Fig. 5c) was quantified as follows:

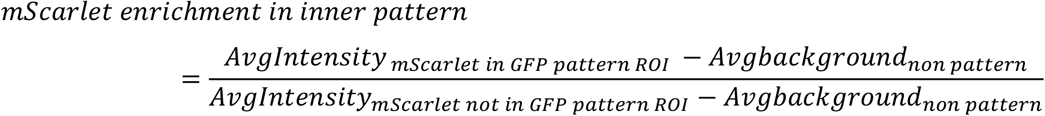

With *AvgIntensity_mScarlet in GFP pattern ROI_* representing the average fluorescence intensity of mScarlet in the Region of interest corresponding to the micropatterned GFP, and *AvgIntensity_mScarlet not in GFP pattern ROI_* representing the average fluorescence Intensity of mScarlet in the Region Of Interest (ROI) of the same size, but corresponding to a ROI in the Fibronectin pattern surrounding the GFP center. *Avgbackground_non pattern_* represents the mScarlet fluorescence intensity in unpatterned regions of the coverslip.

The enrichment of EGFR, and Clathrin (Fig. 6c, f) were calculated as follows:

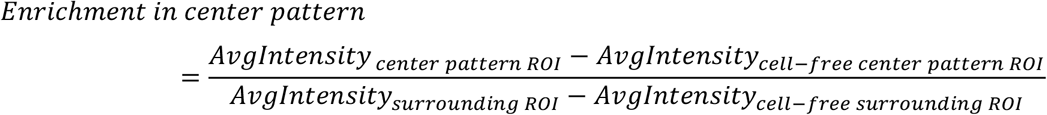

With *AvgIntensity_center pattern ROI_* representing the average fluorescence intensity of EGFR/Clathrin in the region of interest corresponding to the micropatterned EGF (where a cell is adhered to the pattern), *AvgIntensity_cell–free center pattern ROI_* representing the fluorescence signal in a neighbouring region of interest corresponding to micropatterned ligand (but without a cell adhered, to account for potential fluorescence bleed-through into the imaging channel), *AvgIntensity_surrounding ROI_* representing the EGFR/Clathrin signal in the Fibronectin pattern surrounding the EGF center (where a cell is adhered), and *AvgIntensity_cell–free surrounding ROI_* representing the fluorescence intensity in a neighbouring Fibronectin pattern without a cell adhered (to again account for any potential fluorescence bleed-through).

The fold enrichment of GFP-Notch1 (Fig. 6f) was calculated from live imaging of U2OS cells landing and spread on micropatterns, 20 minutes after landing on micropatterns, as follows:

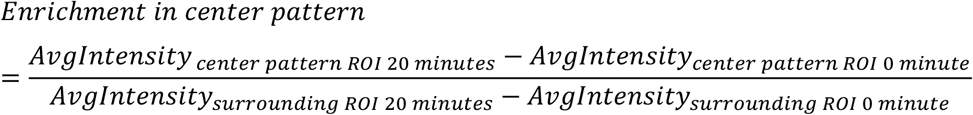

Where *AvgIntensity_center pattern ROI 20 minutes_* is the average fluorescence intensity of GFP-Notch1 20 minutes after a cell landing on the micropattern in the region corresponding to the micropatterned DLL4, and *AvgIntensity_center pattern ROI 0 minutes_* representing the average fluorescence intensity in the same region prior to the cell landing on the micropattern (to account for any potential fluorescence bleed through). Similarly, *AvgIntensity_surrounding ROI 20 minutes_* is the average fluorescence intensity of GFP-Notch1 in the outer micropattern, 20 minutes after a cell lands on the micropattern, and *AvgIntensity_surrounding ROI 0 minutes_* is the average fluorescence intensity of the same region prior to a cell landing on it.

To quantify the recruitment of GFP-Notch1 to DLL4 micropatterns over time (Fig. S8b), the raw increase in GFP-Notch1 signal overlaying the central, DLL4 region compared to the outer fibronectin pattern was calculated (due to relatively low signals, which make calculating the fold change volatile):

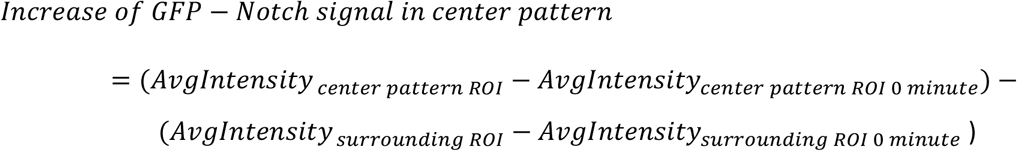

Where *AvgIntensity_center pattern ROI_* is the average fluorescence intensity of GFP-Notch1 in the central, DLL4 pattern at the specific time point, and *AvgIntensity_center pattern ROI 0 minutes_* represents the average fluorescence intensity in the central region prior to a cell landing (to account for any potential fluorescence bleed through). *AvgIntensity_surrounding ROI_* is the average fluorescence intensity of GFP-Notch1 in the outer pattern at the same time point, and *AvgIntensity_surrounding ROI 0 minutes_* is the average fluorescence intensity in the same region, prior to a cell landing on the micropattern (to account for potential bleed through).

For analysis of microtubule gliding on Kin1-biotin patterns (Fig. 3), movies were projected in time, and these projections were used to define the path of gliding microtubules, from which kymographs were generated and analysed using the ‘Kymo Toolbox’ plugin for imageJ, developed by Fabrice Cordelières^37^. For analysis of the proportion of microtubules undergoing directional movement, microtubules were manually segmented based on whether or not they showed clear, directed motion.

To quantify the increased patterning efficiency of S2 cells onto small patterns in Figure 4, we computed the normalized density of micropatterned cells as follows:

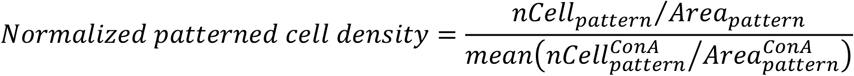

With *nCell_pattern_* representing the number of cells per pattern and *Area_pattern_* representing the area of the same patterns, and 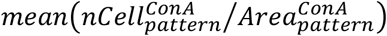 mean density of the ConA sample. Only small patterns, of diameter 20-40 μm were considered for analysis.

To quantify the specificity of S2 cell adhesion, we computed the normalised non-specific adhesion (to patterns of between 20 and 40 μm in diameter) as follows:

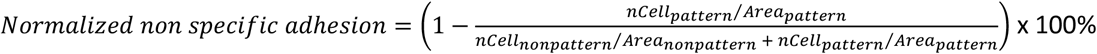

### Statistics

Unless stated otherwise, measurements are given in mean ± SEM. Statistical analyses were performed using GraphPad Prism 8 or SigmaStat 3.5 with an alpha of 0.05. Normality of variables was verified with Kolmogorov-Smirnov tests. Homoscedasticity of variables was always verified when conducting parametric tests. Post-hoc tests are indicated in their respective figure legends.

## Acknowledgments

This work has been supported by the Medical Research Council (MC_UP_1201/13 to E.D) and the Human Frontier Science Program (CDA to E.D.). We thank Vicente Jose Planelles Herrero for help with graphics, and Nicolas Chiaruttini, Jerome Boulanger, James Manton, Christopher Rowlands and Clemens Kaminski for sharing their expertise in optics and microscope design. We thank Manuel Thery and Benoit Vianay for their extensive help to set up micropatterning in the lab at the beginning of this project. We thank Laurent Blanchoin and Antoine Jegou for critically reading the manuscript. We thank the electronics workshop of the LMB, in particular Martin Kyte for building custom LED drivers for this project. We thank the technical instrumentation workshop of the LMB, in particular Steve Scotcher and Adam Fowle for their help in designing and manufacturing the custom-made optical mounts of the DMD-illuminator. We thank Tino Pleiner and Mark Allen for plasmids.

## Author contributions

J.L.W performed all micropatterning experiments, expect Fig. S4, which was performed by S.A. J.L.W purified all proteins and Fibrinogen-anchors, with some assistance from E.D. E.D. developed the LED illuminator with assistance from J.L.W and S.A. B.O.S developed the one-step quantitative synthesis of the BBTB with support from J.C.. A.A.D and SCB provided the purified DLL4 protein and derived the GFP-Notch U2OS cell line. J.L.W, S.A and E.D. produced the figures. E.D. wrote the manuscript. All authors discussed the results and commented on the manuscript.

## Supplementary figures

**Figure S1.**
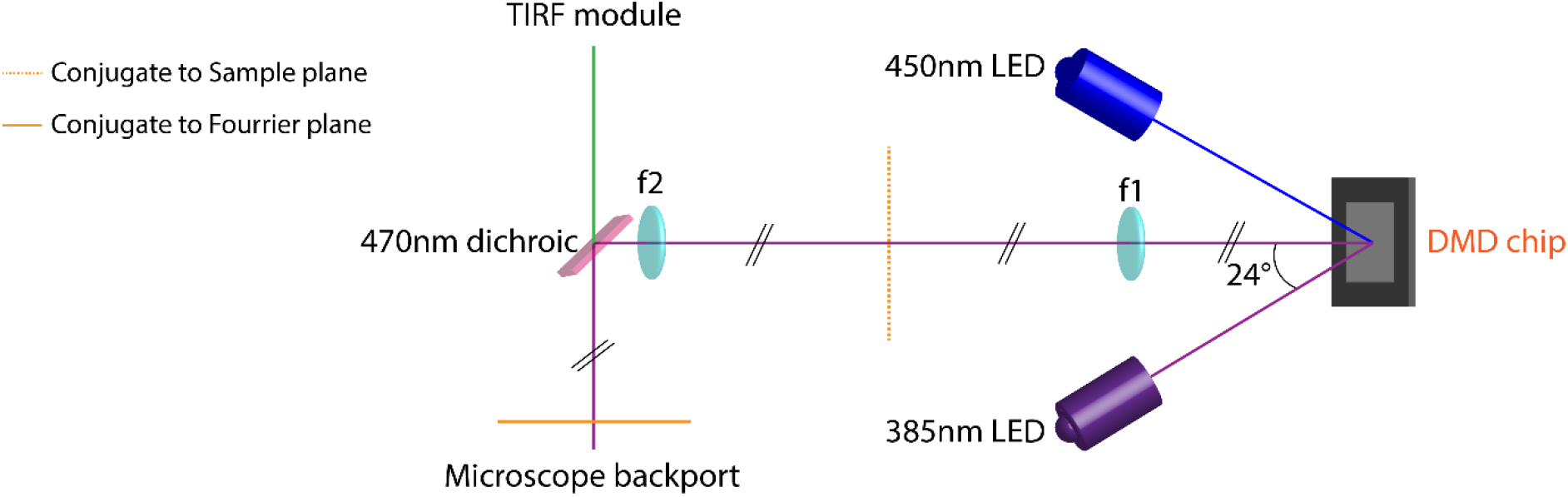
Optical design of the DMD-UV illuminator used in this study. Schematic optical path. A 385 nm high power UV LED light source is collimated using an AR coated aspheric lens and the collimated UV beam is then directed towards a Digital Micromirror Device chip (DMD) at a 24° angle of incidence (corresponding to twice the tilting angle of the DMD mirrors). The image of the DMD chip is then relayed onto the conjugate of the sample plane at the backport of the microscope through a 4f imaging system (f1 = f2 = 125 mm UV Fused Silica Bi-Convex Lenses, AR-Coated). This intermediate image is then relayed onto the sample plane by a tube lens and the objective. To combine DMD-UV illumination with TIRF-illumination, a 470nm dichroic is placed after f2 within a custom backport assembly (Cairn). To offer a second illumination wavelength for 450nm optogenetic stimulation, our design also contains a second 450nm collimated LED at the symmetric −24° angle. This LED can be exchanged for any other LED to provide epifluorescence imaging.

**Figure S2.**
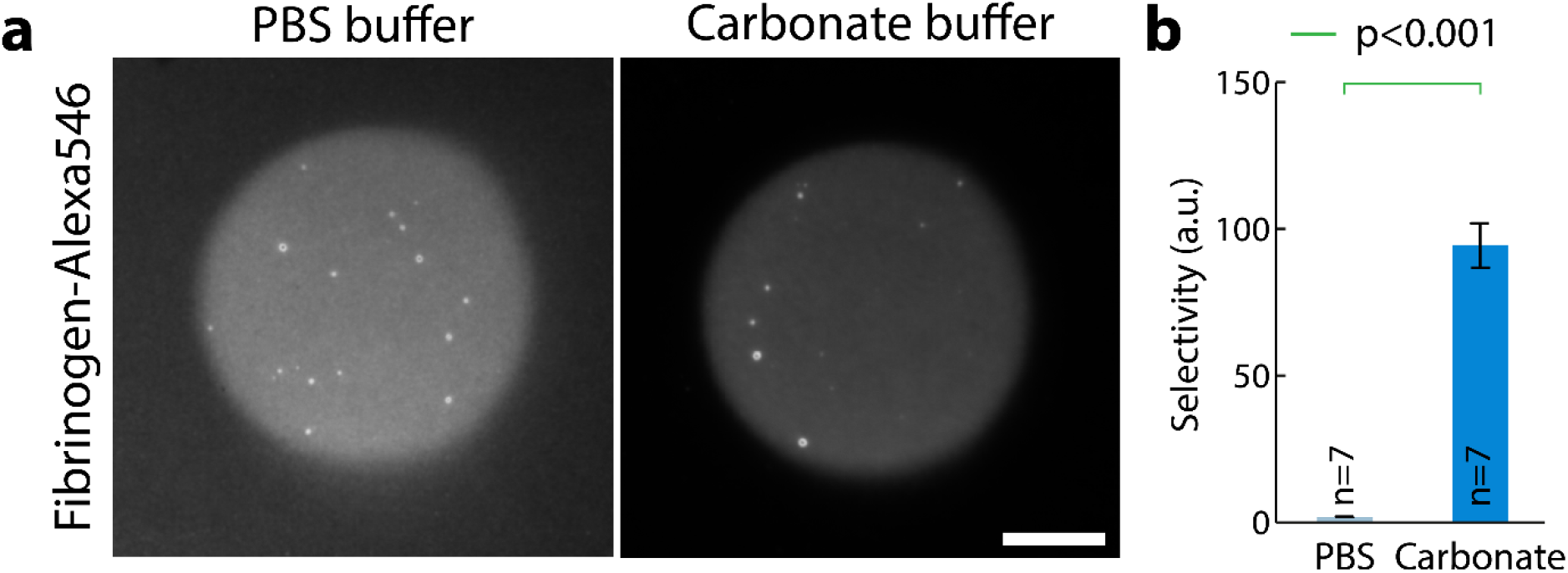
Enhanced Fibrinogen patterning efficiency depends on the buffer used. (a) PLL-PEG-coated glass was processed for LIMAP-patterning with identical UV exposure, pattern and photoinitiator concentration (50 mM BBTB in 0.1 M sodium bicarbonate pH 8.3). Fibrinogen-Alexa546 (50 μg/ml) was then adsorbed onto the UV-activated surface either in PBS or carbonate buffer. After washing, red fluorescence of the patterns was imaged by TIRFM using identical settings (b) Quantification of the effects seen in (a) (average selectivity ± SEM, see methods). Fibrinogen quantitatively patterns better in carbonate buffer. Statistics were performed using a Mann-Whitney Rank Sum Test, n: number of patterns measured. Scale bar: 10μm.

**Figure S3.**
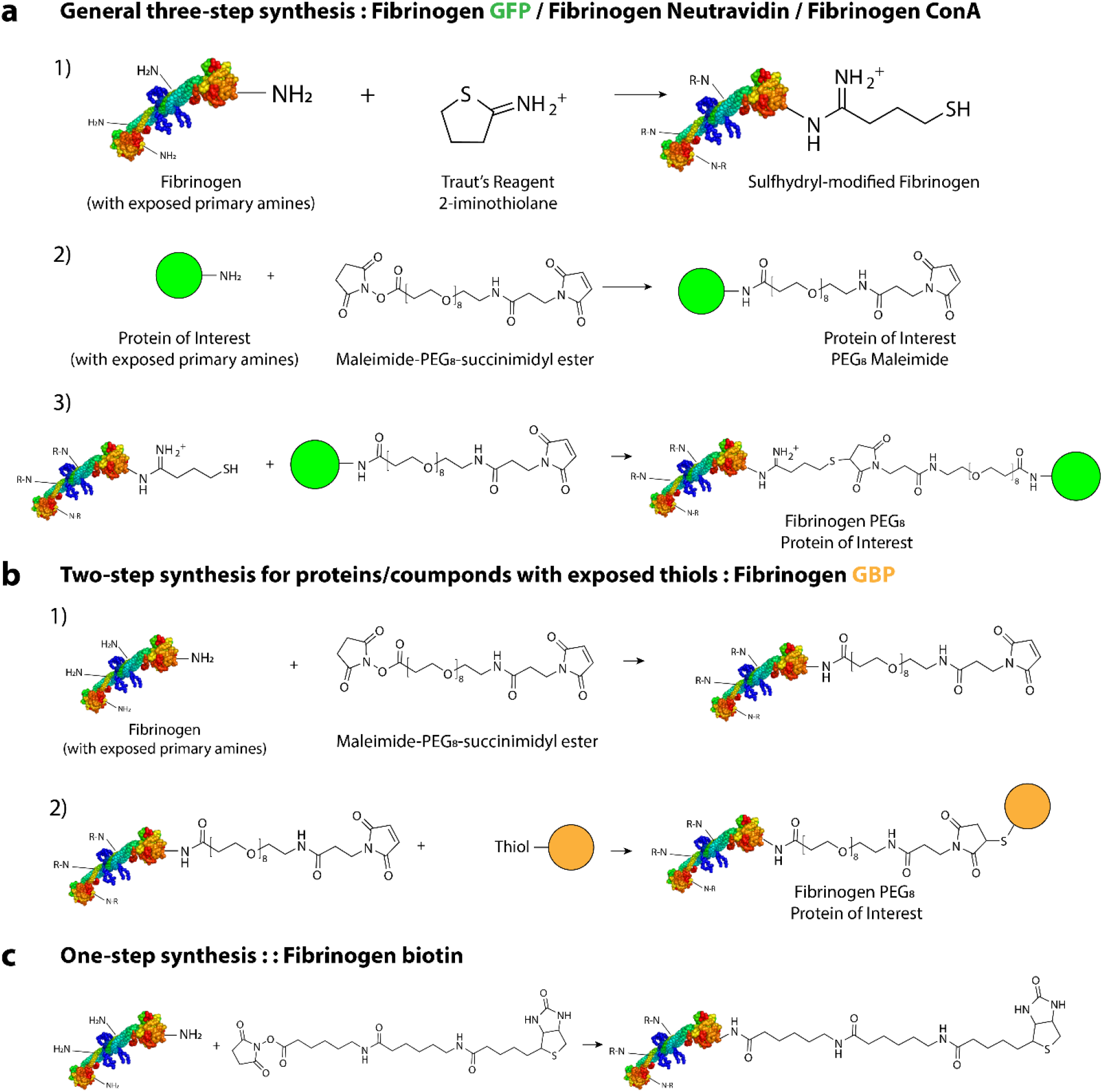
Fibrinoogen is readily functionalized with target proteins or ligands. (a) General three step method to functionalize Fibrinogen with a protein of interest using Lysine reactivity. Reactive thiols are generated onto Fibrinogen, which then are reacted with Maleimide-functionalized protein of interest (GFP, ConA, NeutrAvidin). Excess unreacted protein of interest is removed differential native precipitation. (b) Two step method to functionalize Fibrinogen if protein/ligand of interest has reactive thiols (for example, nanobodies engineered with cysteines). (c) One step method to functionalize Fibrinogen with an NHS derivative of a ligand of interest (Biotin, Fluorophores).

**Figure S4.**
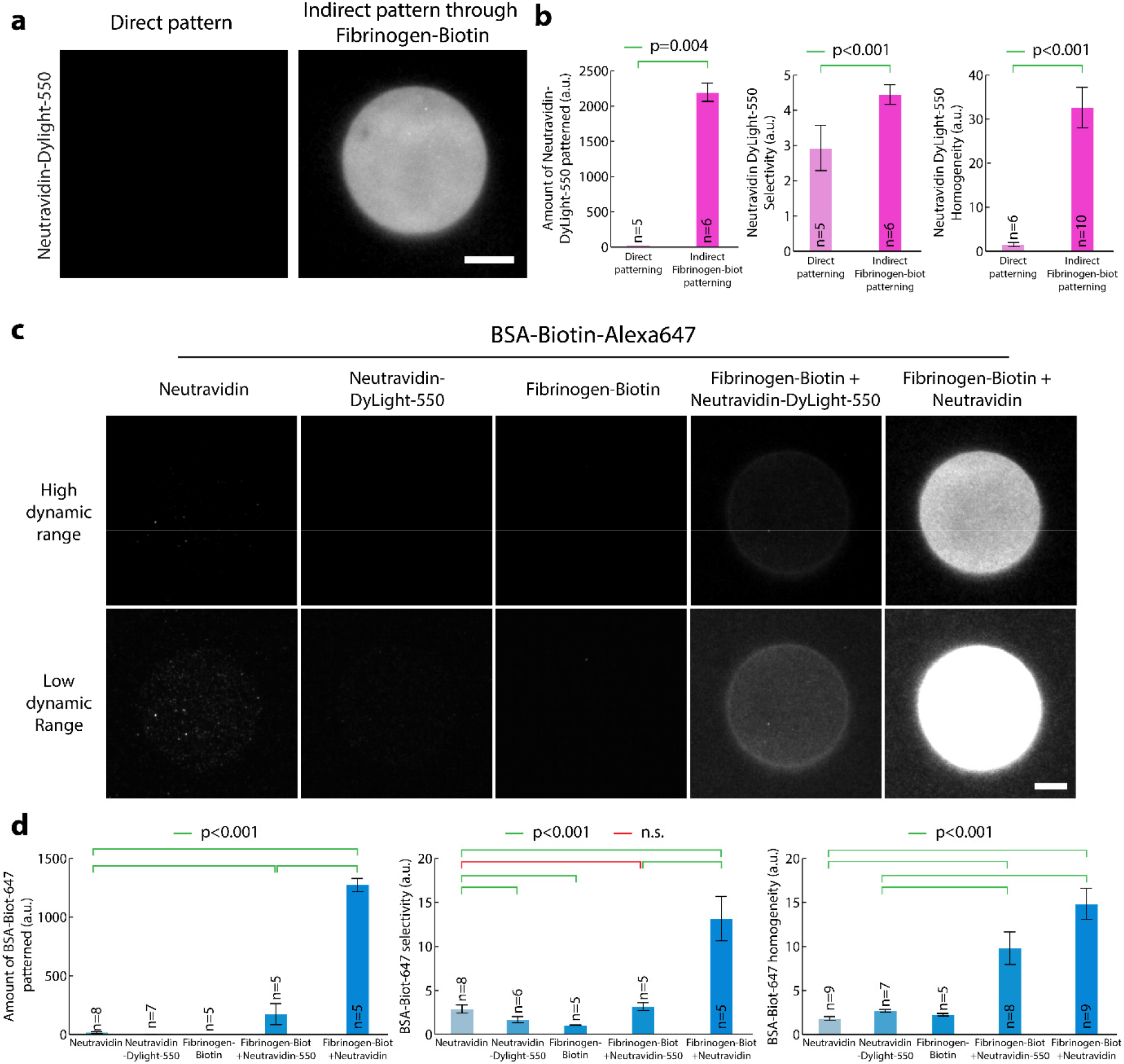
Fibrinogen anchors improves selectivity and homogeneity of micropatterns on PEG-Silane surfaces. (a) NeutrAvidin-DyLight-550 (50 μg/ml) was micropatterned on PLL-PEG-coated glass using LIMAP. Alternatively, Fibrinogen-biotin was micropatterned with identical UV exposure, pattern shape and protein concentration, followed by the addition of NeutrAvidin-Dylight-550 and imaging by TIRFM. (b) Quantification of pattern selectivity (left panel), amount of micropatterned protein (middle panel) and homogeneity (right panel) in the sample presented in (a). Statistics were performed using a Mann-Whitney test for the left and right panels, and a Student’s t-test for the middle panel. Fibrinogen biotin significantly improves the patterning of NeutrAvidin-Dylight-550. (c) NeutrAvidin, NeutrAvidin-Dylight-550 or Fibrinogen-biotin were micropatterned on PLL-PEG-coated glass using LIMAP with identical UV exposure, pattern shape and protein concentration (50 μg/ml). After pattern quenching and washing, BSA-Biotin-Alexa647 (5 μg/ml) was added for 5 min, then the sample was washed and BSA-Biotin-Alexa647 fluorescence was imaged by TIRFM (NeutrAvidin, NeutrAvidin-Dylight-550 and Fibrinogen-biotin samples). Alternatively, Fibrinogen-biotin was micropatterned as above, then NeutrAvidin (or NeutrAvidin-Dylight-550) was added before addition of BSA-Biotin-Alexa647 and TIRFM imaging. Two different exposures were used for each sample (top versus bottom line) so that each lane could be represented with the same dynamic range. (d) Quantification of pattern selectivity (left panel), amount of micropatterned protein (middle panel) and homogeneity (right panel) in the sample presented in (c). Statistics were performed using an ANOVA1 test followed by a Tukey post-hoc test after log10 transformation of the data (p<0.001). Fibrinogen-biotin enhances significantly the amount of BSA-Biotin-Alexa647 patterned, as well as pattern selectivity and the homogeneity. While Fibrinogen biotin efficiently improves the micropatterning of NeutrAvidin-Dylight-550 compared to direct micropatterning, this does not translate into improved micropatterning of BSA-Biotin-Alexa647, likely because NeutrAvidin-Dylight-550 has lost some biotin-binding activity due to its fluorescent labelling. Scale bar: 10μm.

**Figure S5.**
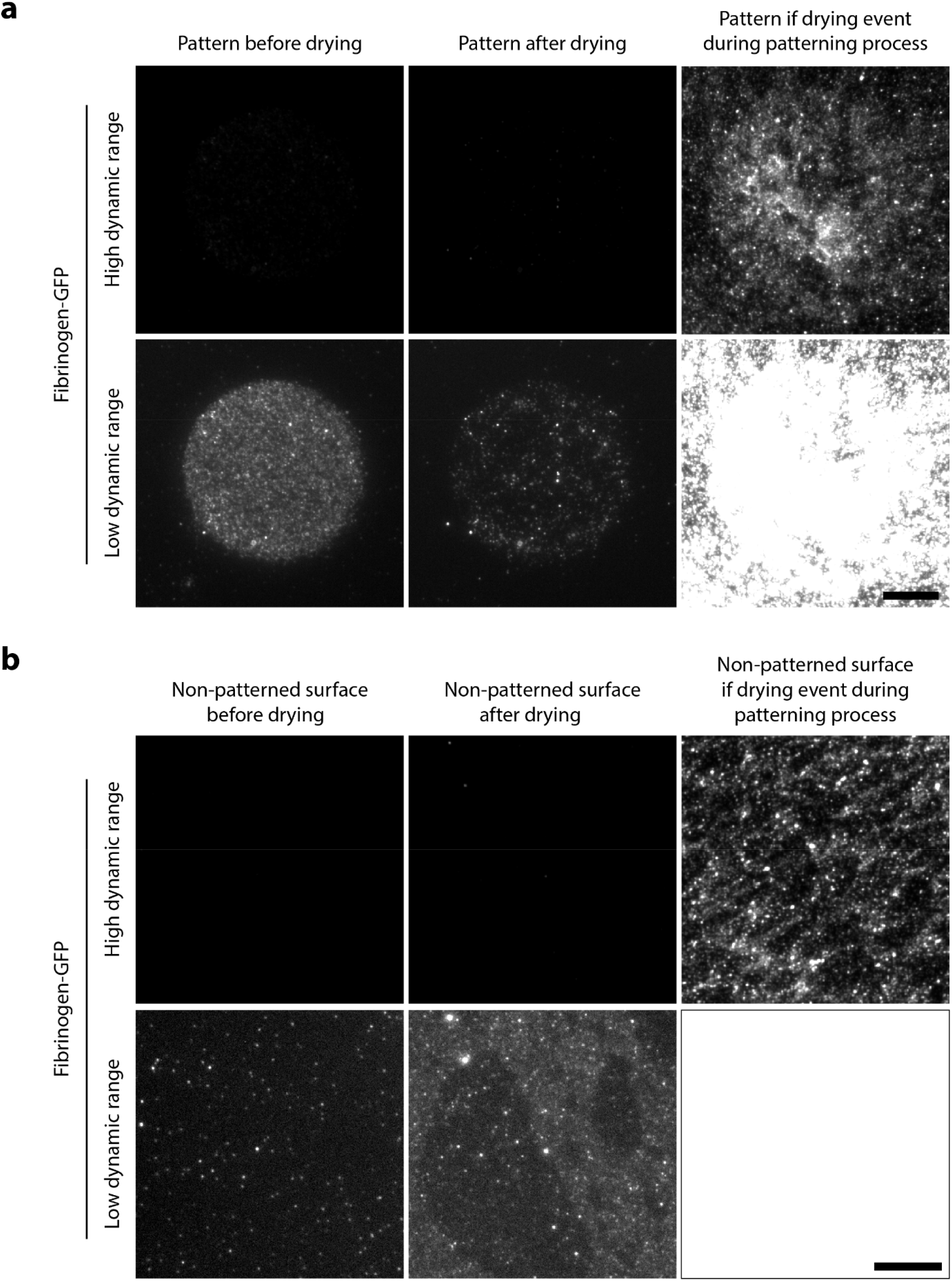
Sample drying during or after the patterning process affects patterning efficiency. (a) Drying of the pattern during or after the patterning process negatively impacts the patterning efficiency. Left panels: PLL-PEG-coated glass was processed for LIMAP-patterning of Fibrinogen-GFP (50 μg/ml). Middle panels: after imaging the pattern, the sample was dried, then rehydrated and imaged again in the same conditions. Right panels: PLL-PEG-coated glass was processed for LIMAP-patterning of Fibrinogen-GFP as before, but sample was allowed to dry after the adsorption process. Two different dynamic range for visualisation were used for each sample (top versus bottom line) so that each lane could be represented with the same dynamic range. (b) Protein adsorption to the nonpatterned region is also increased upon drying during/after the patterning process. Sample was treated as in A, but non-patterned regions were imaged. Scale bar: 10 μm.

**Figure S6.**
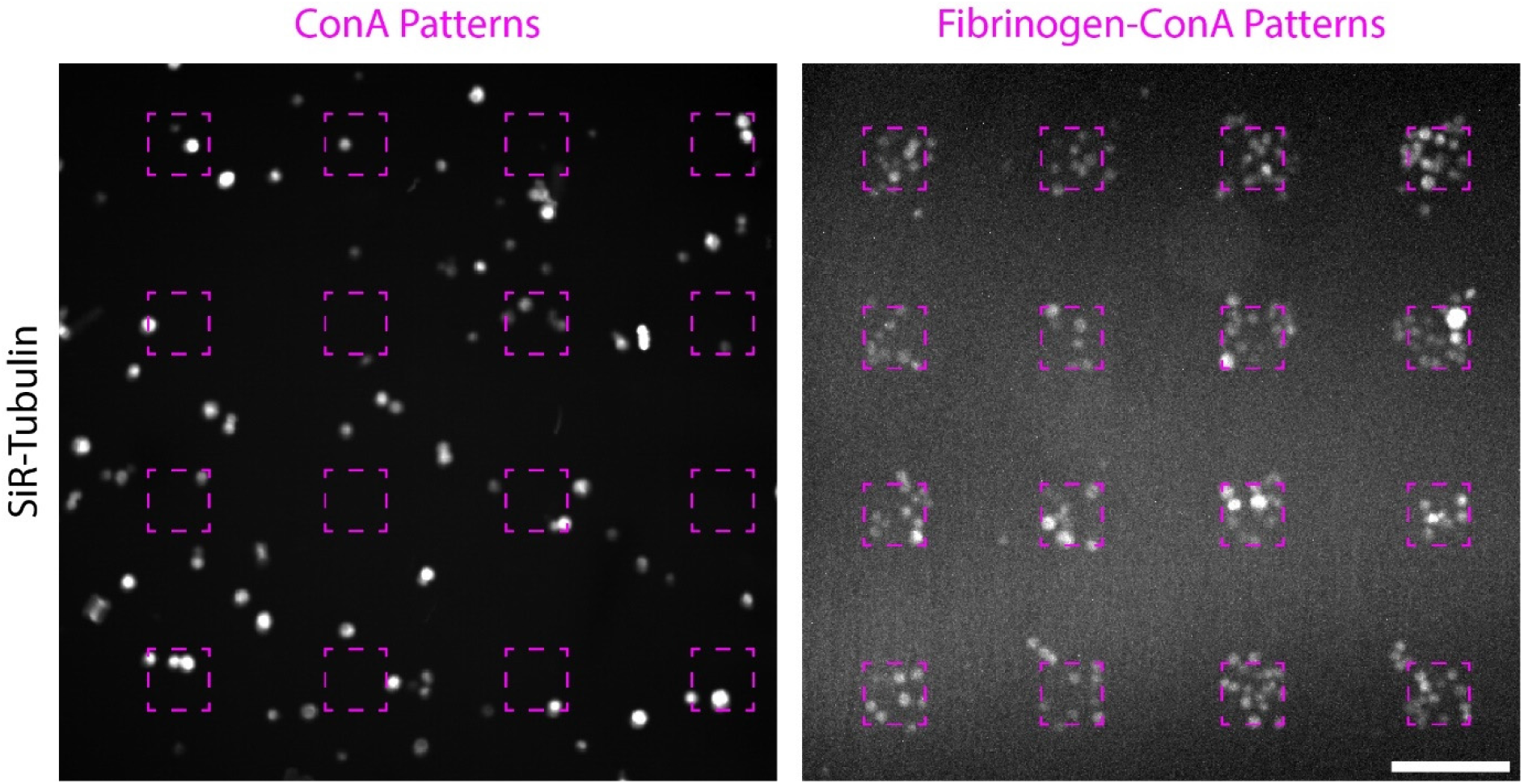
Fibrinogen anchors enahnces the micropatterning of hard-to-pattern cells. ConA (doped with 10% Rhodamine ConA) or Fibrinogen-ConA (doped with 10% Fibrinogen-Alexa546) were micropatterned at 50 μg/ml onto PLL-PEG-coated glass using deep-UV and a chromium mask. Coverslips were washed, and S2 cells were added for 1h, before addition of SiR tubulin, to label cells, for 30 minutes. After washing, cells and micropatterns were imaged by spinning disc confocal microscopy. Micropatterning efficiency of S2 cells is lower on ConA compared to Fibrinogen-ConA. Dashed line: region exposed to deep-UV. Scale bar: 100 μm.

**Figure S7.**
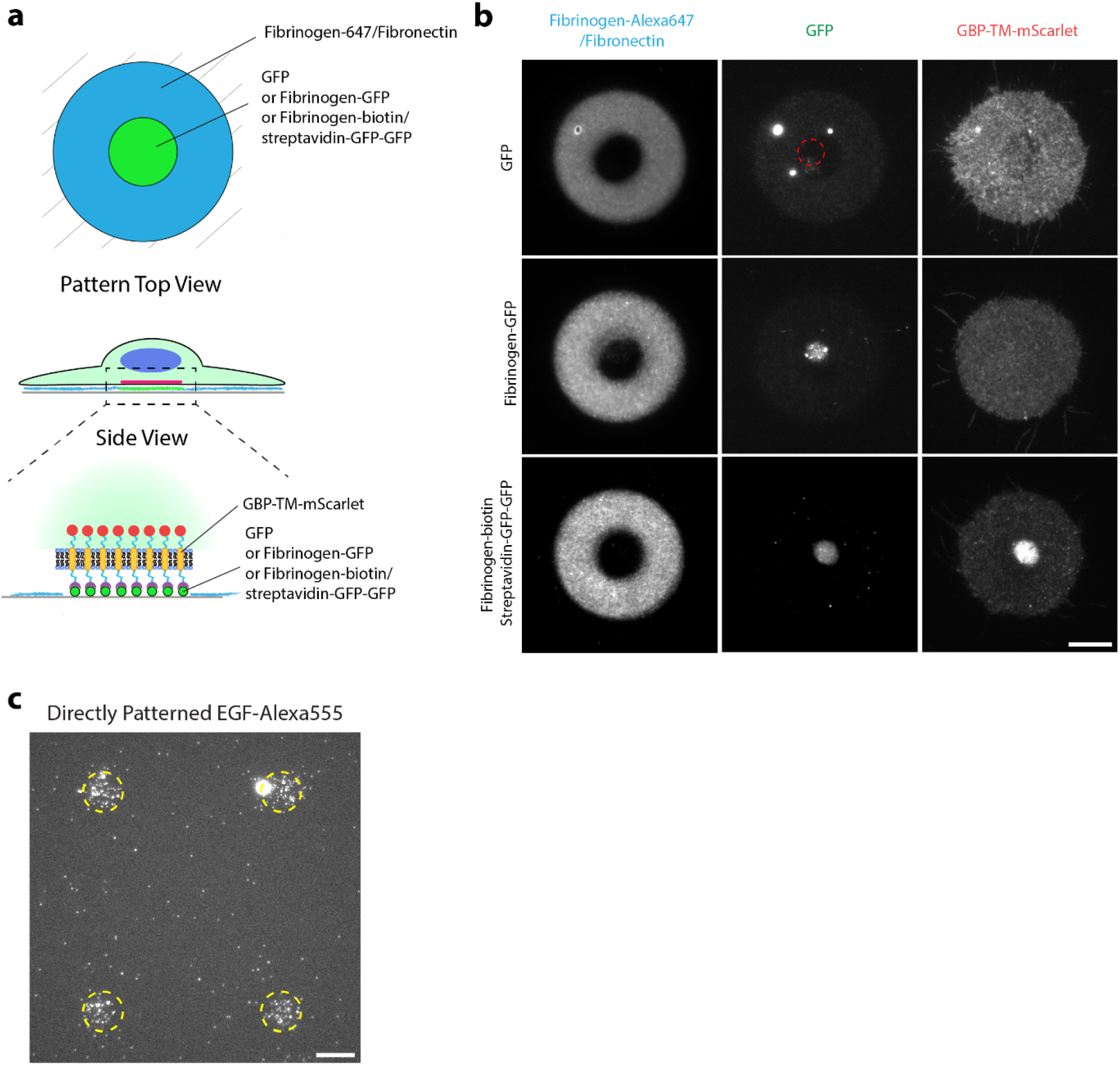
Regular micropatterning does not allow subcellular micropatterning of receptors. (a) Experimental scheme: stable NIH/3T3 cells constitutively expressing GBP-TM-mScarlet were allowed to spread on dual patterns of Fibronectin/Fibrinogen-Alexa647 and either GFP, or Fibrinogen-GFP (low degree of labelling of 0.5 mol GFP per mol fibrinogen), or Fibrinogen-biotin/streptavidin-GFP-GFP then imaged live by TIRF confocal microscopy. (b) Only high-density GFP micropatterning via Fibrinogen-biotin/streptavidin-GFP-GFP allows efficient relocalisation of the GBP-TM-mScarlet construct onto an area defined by the extracellular pattern. Note that bottom panel corresponds to Fig. 5b, reproduced here for convenience. Note also that GFP channels are not intensity-matched here. In reality, there is much more GFP when using streptavidin-GFP-GFP as compared to when using Fibrinogen-GFP. (c) In contrast to when biotin-EGF was attached to the Fibrinogen-Biotin-ATTO490LS/NeutrAvidin sandwich (Fig. 6a-c), direct patterning of biotin-EGF/streptavidin-Alexa555 (1 μg/ml) showed very weak, inhomogeneous and non-specific patterning, preventing patterning of the second, surrounding Fibronectin pattern. Dashed line: region exposed to UV. Scale bars: 10 μm.

**Figure S8.**
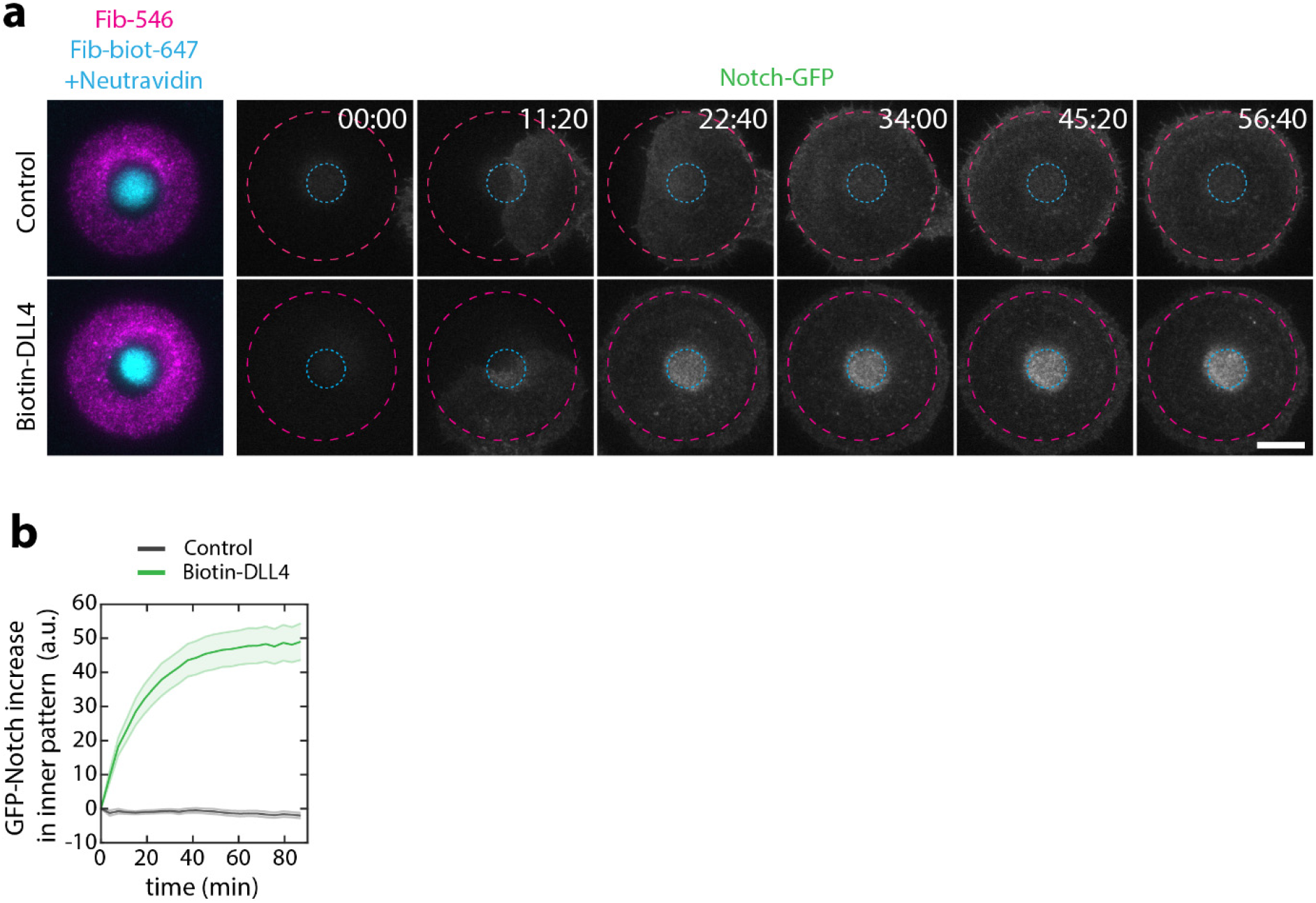
Dynamics of GFP-Notch relocalisation by Delta micropatterns. (a) Experimental scheme: U2OS cells stably expressing GFP-Notch1 were allowed to spread on dual patterns of Fibronectin/Fibrinogen-Alexa647 and Fibrinogen-Biotin-ATTO490LS/NeutrAvidin/biotin-DLL4 (see Fig 6d,e) and GFP-Notch fluorescence was imaged live during spreading by TIRF microscopy. Elapsed time in min:sec. (b) Quantification of the effects seen in (a), see also methods. Scale bar: 10 μm.

## Supplementary movie legends

**Movie S1 Fibrinogen anchors facilitate micropatterning of active motors.**

Biotinylated-Kinesin1 motors (Kin1-Biotin) were micropatterned on PLL-PEG-coated glass using LIMAP either directly or indirectly through a Fibrinogen-Biotin-ATTO490LS/NeutrAvidin sandwich. After washing and quenching, GMPCPP-stabilized fluorescent microtubules were added in the presence of ATP and their motion observed by TIRFM. Dashed purple line delineate the kinesin pattern as imaged either through post-labelling of Kin1-Biotin by streptavidin-Alexa647 for direct patterning or by Fibrinogen-Biotin-ATTO490LS fluorescence for the indirect labelling. This movie corresponds to Fig. 3. Scale bars: 10 μm

**Movie S2 Dynamics of GBP-TM-mScarlet relocalisation by Fibrinogen-GFP micropatterns**

NIH/3T3 cells stably and constitutively expressing GBP-TM-mScarlet were allowed to spread on dual patterns of Fibronectin/Fibrinogen-Alexa647 and Fibrinogen-Biotin-ATTO490LS/streptavidin-GFP-GFP, and were then imaged live by TIRF microscopy. Elapsed time in min:sec. This movie corresponds to Fig. 5d. Scale bar: 10 μm.

**Movie S3 Dynamics of GFP-Notch relocalisation by Fibrinogen-Delta micropatterns.**

U2OS cells stably expressing GFP-Notch1 were allowed to spread on dual patterns of Fibronectin/Fibrinogen-Alexa647 and Fibrinogen-Biotin-ATTO490LS/NeutrAvidin/biotin-DLL4 (see Fig 6d,e) and GFP-Notch fluorescence was imaged live during spreading by TIRF microscopy. Elapsed time in min:sec. This movie corresponds to Fig. S8a. Scale bar: 10 μm.

## Bibliography

1. Bumgarner, R. Curr. Protoc. Mol. Biol. (2013) doi:10.1002/0471142727.mb2201s101.

2. Braunschweig, A. B., Huo, F. & Mirkin, C. A. Nat. Chem. (2009) doi:10.1038/nchem.258.

3. Falconnet, D., Csucs, G., Michelle Grandin, H. & Textor, M. Biomaterials (2006) doi:10.1016/j.biomaterials.2005.12.024.

4. Thé, M. et al. Proc. Natl. Acad. Sci. U. S. A. (2006) doi:10.1073/pnas.0609267103.

5. Théry, M., Jiménez-Dalmaroni, A., Racine, V., Bornens, M. & Jülicher, F. Nature (2007) doi:10.1038/nature05786.

6. Reymann, A. C. et al. Science (80-.). (2012) doi:10.1126/science.1221708.

7. Schaedel, L. et al. Nat. Phys. (2019) doi:10.1038/s41567-019-0542-4.

8. Aumeier, C. et al. Nat. Cell Biol. (2016) doi:10.1038/ncb3406.

9. Toro-Nahuelpan, M. et al. Nat. Methods (2020) doi:10.1038/s41592-019-0630-5.

10. Engel, L. et al. J. Micromechanics Microengineering (2019) doi:10.1088/1361-6439/ab419a.

11. Azioune, A., Storch, M., Bornens, M., Théry, M. & Piel, M. Lab Chip (2009) doi:10.1039/b821581m.

12. Waldbaur, A., Waterkotte, B., Schmitz, K. & Rapp, B. E. Small (2012) doi:10.1002/smll.201102163.

13. Bélisle, J. M., Kunik, D. & Costantino, S. Lab Chip (2009) doi:10.1039/b911967a.

14. Eichinger, C. D., Hsiao, T. W. & Hlady, V. Langmuir (2012) doi:10.1021/la2039202.

15. Fink, J. et al. Lab on a Chip (2007) doi:10.1039/b618545b.

16. Strale, P. O. et al. Adv. Mater. (2016) doi:10.1002/adma.201504154.

17. Portran, D., Gaillard, J., Vantard, M. & Thery, M. Cytoskeleton (2013).

18. Godinho, S. A. et al. Nature (2014) doi:10.1038/nature13277.

19. Carpi, N., Carpi, N., Piel, M., Azioune, A. & Fink, J. Protoc. Exch. (2011) doi:10.1038/protex.2011.238.

20. Rogers, S. L., Rogers, G. C., Sharp, D. J. & Vale, R. D. J. Cell Biol. (2002) doi:10.1083/jcb.200202032.

21. Rothbauer, U. et al. Mol Cell Proteomics (2008).

22. Bachand, G. D., Bouxsein, N. F., Vandelinder, V. & Bachand, M. Wiley Interdiscip. Rev. Nanomedicine Nanobiotechnology (2014) doi:10.1002/wnan.l252.

23. Subramanian, R. & Gelles, J. J. Gen. Physiol. (2007) doi:10.1085/jgp.200709866.

24. Pleiner, T., Bates, M. & Görlich, D. J. Cell Biol. (2018) doi:10.1083/jcb.201709115.

25. Pleiner, T. et al. Elife (2015).

26. Schmidt, T. G. M. & Skerra, A. Nat. Protoc. (2007) doi:10.1038/nprot.2007.209.

27. Veggiani, G. et al. Proc. Natl. Acad. Sci. U. S. A. (2016) doi:10.1073/pnas.1519214113.

28. Yamaguchi, H., Chang, S. S., Hsu, J. L. & Hung, M. C. Oncogene (2014) doi:10.1038/onc.2013.74.

29. Derivery, E. et al. Nature (2015).

30. Watson, J. L., Stangherlin, A. & Derivery, E. Methods Mol. Biol. (2020) doi:10.1007/978-1-0716-0463-2_10.

31. Bindels, D. S. et al. Nat. Methods (2016) doi:10.1038/nmeth.4074.

32. Derivery, E. et al. Dev Cell (2009).

33. Malecki, M. J. et al. Mol. Cell. Biol. (2006) doi:10.1128/mcb.01655-05.

34. Hueck, K. (2016) doi:10.5281/ZENODO.47513.

35. Lukinavicius, G. et al. Nat Methods (2014).

36. Schindelin, J. et al. Nat. Methods (2012).

37. Zala, D. et al. Cell (2013) doi:10.1016/j.cell.2012.12.029.

